# Hydrophobic and lipid-mediated gating mechanism revealed by low-conductance MthK mutants

**DOI:** 10.64898/2026.04.28.721335

**Authors:** Reinier de Vries, Hee-Seop Yoo, Chen Fan, Wojciech Kopec, Crina M. Nimigean, Bert L. de Groot

## Abstract

Large-conductance calcium (Ca^2+^)-activated potassium (K^+^) channels are involved in several essential cellular processes. This requires the ability to alternate between conductive and non-conductive states, yet the molecular mechanism of their gating remains incompletely understood. Bundle-crossing, selectivity filter, and hydrophobic gating are some of the mechanisms previously proposed for these channels, including the archaeal Ca^2+^-activated K^+^ channel MthK. Here, we took advantage of two MthK mutants that show dramatically-reduced conductance to gain insights into how the channels gate at the molecular level by combining cryo-electron microscopy, functional assays, and extensive molecular dynamics simulations. Functional and computational analyses reveal distinct mechanisms of reduced conductance between the two mutants. One mutant, where a glutamate residing at the intracellular mouth of the pore was replaced by an alanine (E92A) reduced ion density in the cavity and dramatically accelerated spontaneous channel closure. The second mutant, where an alanine residing in the channel’s aqueous cavity below the selectivity filter was replaced by a phenylalanine (A88F), introduced a hydrophobic constriction that partially dehydrates the cavity, imposing a barrier to ion permeation. The cryo-EM structures of these mutants, virtually identical to those of WT MthK, were used as models to investigate gating using molecular dynamics (MD) simulations. Long-timescale simulations captured complete open-to-closed transitions in both mutants, uncovering a dehydrated intermediate state and a tightly coupled lipid-entry mechanism that drove final pore closure. Our results show that hydrophobicity and lipid interactions are central determinants of MthK gating.

## Introduction

Potassium channels are the most widely distributed type of ion channels and allow for selective permeation of potassium ions across cell membranes^1,2^, which is essential for a variety of biological processes. The ability of these channels to change between permeating and non-permeating states is of vital importance to regulate these processes. While the overall structure and especially the selectivity filter (SF) of potassium channels is highly conserved^3,4^, various different mechanisms have evolved to stop ion permeation in these channels. Gating and inactivation can occur at different parts of the channel pore, such as the intracellular bundle-crossing, the selectivity filter, and other constrictions in the aqueous cavity^3,5–8^. A different mechanism from pore constriction is hydrophobic gating, where dehydration of the pore cavity can result in an impermeable barrier without a sterically closed lower gate^9–11^.

MthK is an archaeal calcium-gated potassium channel and has been investigated as a model potassium channel^8,12^ due to its similarity to the eukaryotic BK channels. The channel consists of a minimal pore domain composed of only two transmembrane helices, and an intracellular, calcium-binding RCK domain. These channels open by binding of intracellular Ca^2+^ ions to the RCK gating ring, which triggers a conformational change that is transmitted through the linker region to the inner (TM2) helices of the pore, leading to channel opening at the inner bundle-crossing gate^8,13,14^.

Several high-resolution structural studies have provided snapshots of MthK in open^5,13,15,16^, closed^15,17^ and inactivated^15^ conformations, yet the precise gating mechanism, that is, how the channel transitions between these states is still unknown. Hydrophobic gating has been proposed as a gating mechanism in two-pore domain potassium K_2_P channels, where dewetting of a hydrophobic pore segment leads to functional closure in the absence of a steric gate^18,19^. This phenomenon has also been suspected to occur in other channels, such as BK channels and MthK^20^, where spontaneous lipid entry through side-facing fenestrations has been observed in molecular dynamics (MD) simulations^21,22^. In addition, no bundle-crossing constriction was detected in BK channels in the absence of Ca^2+^, suggesting that the BK cavity is accessible in both open and closed states^20,23–26^. On the other hand, cryoEM studies of closed MthK have revealed a helix bundle-crossing constriction at the intracellular mouth of the pore, similar to KcsA^27,28^, which forms a steric block for ion permeation in this closed state^15^. Nevertheless, recent MD simulations^20^ supported by evidence of functional data from hydrophobic MthK mutants at the lower gate and in the pore cavity^29,30^, suggest that MthK may also employ hydrophobic gating under certain conditions.

One MthK residue of particular interest in this context is a glutamate located at the intracellular gate of the pore (E92). Single channel electrophysiology recordings show that mutation of this residue to alanine to make it more hydrophobic can lead to drastically reduced open probability with little reduction in single channel current^30^. Another residue of interest is an alanine located in the aqueous pore cavity (A88) which was mutated to a phenylalanine to increase the hydrophobicity of the cavity. However, single-channel recording data showed that A88F reduces current by a different mechanism from E92A: it displays reduced single-channel conductance with little reduction in open probability^29,30^.

In this work, we combine Cryo-EM, functional data, and atomistic MD simulations to investigate the gating mechanism of MthK, with a particular emphasis on the role of cavity hydration, lipid access, and the molecular determinants of hydrophobic gating. To optimize conditions for detecting hydrophobic gating in MthK we take advantage of the two MthK mutants that increase the hydrophobicity in the cavity: E92A and A88F. Cryo-EM structures of E92A in the presence of Ca^2+^ revealed open conformations only, identical to those found in WT, despite the very low activity previously reported for this mutant. This finding pointed towards a potential hydrophobic gating mechanism where permeation was arrested without steric bundle-crossing closure. Extensive MD simulations revealed that the activity reduction in E92A was indeed due to cavity dehydration followed by bundle-crossing closure. Conversely, the reduction in single-channel conductance in A88F was due to an increased barrier for ion permeation due to the increased cavity hydrophobicity. In addition, long MD simulations reveal the complete transition process in the E92A mutant and reveal a previously unknown intermediate state and highlight the role lipids play in this gating mechanism. We show that cavity dehydration and subsequent lipid entry are the two key steps that define the open-closed transition in MthK.

## Results and Discussion

### Structures of E92A MthK reveal an open-like conformation in the presence of Ca^**2+**^

The previous reports^30^ of extremely low open probability of E92A MthK, raising the expectation that cryo-EM structures of these mutant channels will display only closed state conformations under both the absence and presence of Ca^2+^ conditions. Contrary to this expectation, cryo-EM analysis of E92A MthK reconstituted into lipid nanodiscs revealed an open bundle-crossing conformation in the presence of Ca^2+^ that closely matches the WT structure, while in the absence of Ca^2+^ the mutant adopts the expected closed conformation (Fig. 1A–C, Fig. S1).

**Figure 1.**
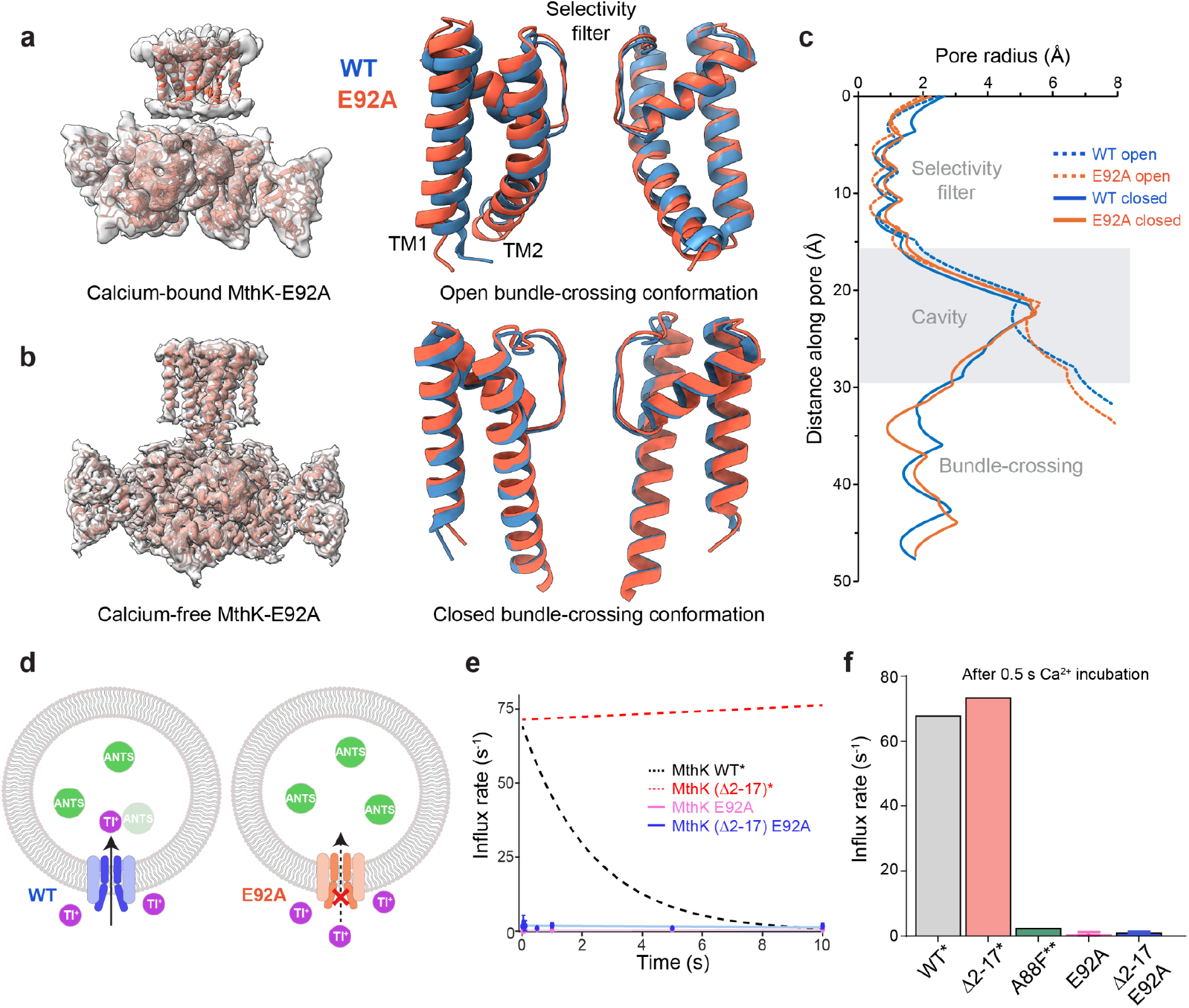
Structural and functional comparison of WT and E92A MthK channels. (**A)** Cryo-EM structure of the MthK E92A mutant in the presence of Ca^2+^ (left) and structural alignment of transmembrane (TM) domain from Ca^2+^-bound WT (PDB: 6U6H; blue) and E92A (orange), showing a similar open bundle-crossing conformation. **(B)** Cryo-EM structure of MthK E92A mutant in the absence of Ca^2+^ (left) and structural alignment of transmembrane domain from Ca^2+^-free WT (PDB: 6U6D; blue) and E92A (orange), indicating similar closed-bundle-crossing conformation. Only two subunits were shown for clarity (A and B). **(C)** Pore radius profiles along the ion conduction pathway for the TM domains of WT and E92A mutant in the open and closed conformations. **(D)** Schematic illustration of the stopped-flow fluorescence assay used to monitor thallium (Tl^+^) influx into proteoliposomes following Ca^2+^ application. The fluorophore ANTS (green), quencher Tl^+^ (purple), MthK WT (blue) and E92A mutant (orange) are depicted. Channels were activated by Ca^2+^, followed by Tl^+^, which quenches the ANTS fluorescence. Fluorescence traces were recorded after varying Ca^2+^ incubation times. **(E)** Tl^+^ influx rates of WT MthK and indicated mutants were plotted as a function of Ca^2+^ incubation time. Data for WT and (Δ2-17) mutants were adapted from previous study.^31^ **(F)** Summary of Tl^+^ influx rates following 0.5 s Ca^2+^ incubation, showing the absence of Ca^2+^-activated flux in E92A MthK. Data for MthK A88F were adapted from a previous study.^17^ Data in panel (E) and (F) represent mean ± s.d. from three independent experiments.

Despite the presence of an open conformation in the structural dataset, E92A MthK proteoliposomes exhibited no response to Ca^2+^ in stopped-flow Tl^+^ influx assays, consistent with previous reports (Fig. 1D and E).^29^ We ruled out fast inactivation as a reason for the lack of activity in these mutants because an inactivation-removed E92A MthK where the N-terminus responsible for inactivation was cleaved off^15^ (Δ2-17 E92A MthK), also showed little activity in stopped-flow assays (Fig. 1E and F). Additional alchemical free energy calculations show that N-type inactivation is less favourable in E92A compared to the WT channel (Fig. S2). Together, these results demonstrate that the absence of Ca^2+^-activated Tl^+^ influx despite an open-like bundle-crossing conformation implies a conduction defect downstream of helix opening, such as pore dewetting and reduced ion occupancy.

### MD simulations of E92A MthK shows decrease in conductance

To investigate the molecular mechanism behind this difference in activity and permeation rate, we performed molecular dynamics simulations of both WT and E92A MthK pores in a POPC bilayer, first focusing on the open conformation we observe that the simulated current is reduced by about 30% in the E92A mutant when compared to the WT channel (Fig. 2B), in agreement with previous single channel recordings in the E92A mutant.^30^ We observe a similar current reduction in full length channels and in control simulations with a different membrane composition.(Fig. S3) .

**Figure 2.**
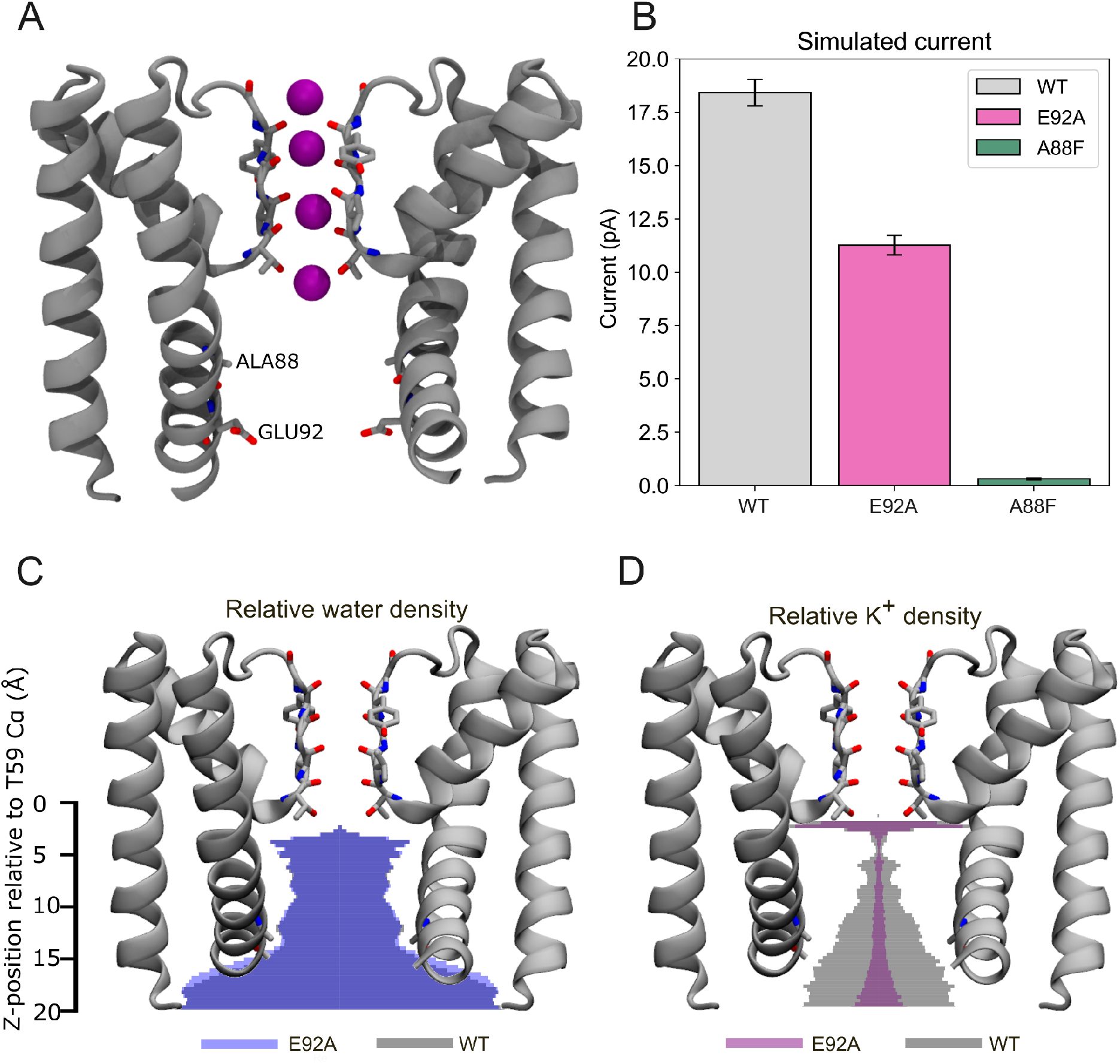
MthK structure and features of more hydrophobic cavity mutants. **(A)** 3D structure of the MthK pore, highlighting A88 and E92, showing just 2 of 4 monomers for clarity. **(B)** Simulated single channel current under 300 mV restrained to the open conformation, the error bar is the SE of the mean (N = 10). **(C)** Simulated water density in the pore cavity relative to the S4 SF binding site in E92A mutant overlaid on the WT density. **(D)** Simulated K^+^ ion density in the pore cavity relative to the S4 SF binding site in E92A mutant overlaid on the WT density.

As the E92A mutation greatly affects the properties of the pore cavity, we looked at the distribution of water molecules (Fig. 2C) and potassium ions (Fig. 2D) in the cavity. We find that that K^+^ density in the whole cavity is greatly reduced in the E92A mutant. Suggesting that this lower effective K+ concentration is the main reason for the observed lower single channel current in the E92A mutant.

### Principal Component Analysis reveals an intermediate state in E92A MthK

In our unrestrained simulations, the E92A mutant became non-conductive after tens to hundreds of nanoseconds. Visual inspection of these trajectories showed that in these simulations the channels spontaneously dehydrate and close. To quantify and track this process we performed a Principal Component Analysis (PCA) on a set of available X-ray and Cryo-EM structures (Fig. S6 & Table S2). The first principal component (PC1) from this analysis allows us to separate and identify open (PC1 ≈ -1 nm) and closed states (PC1 ≈ 11 nm). Free simulations of the WT MthK pore domain started from the open and closed state, stay near the initial state and are clearly separated along PC1 (Fig. 3A-B). For the E92A mutant, however, the open-closed intermediate region is also sampled (PC1 ≈ 5 nm) (Fig. 3C), confirming partial closing of the channel.

**Figure 3.**
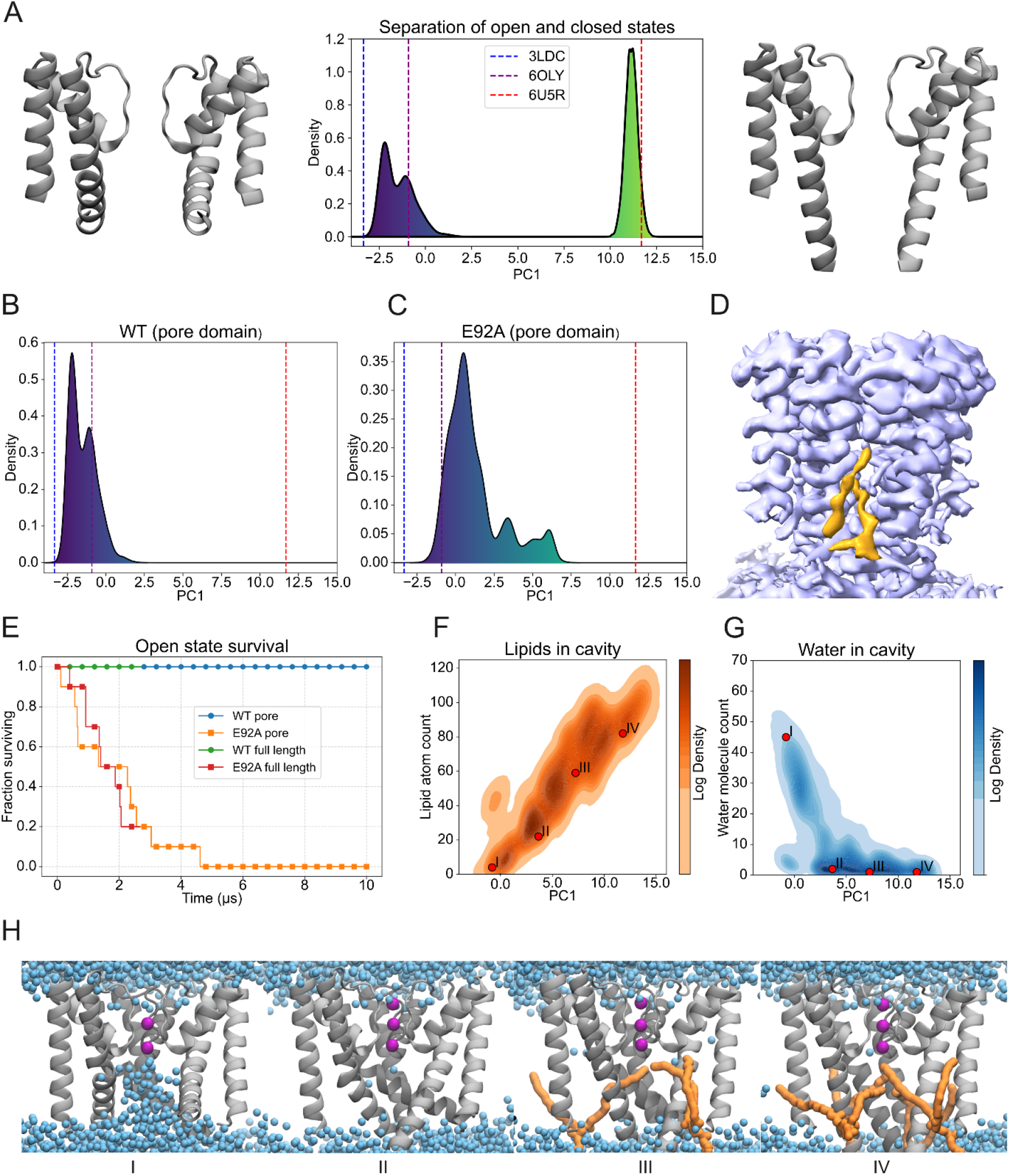
MD simulations of MthK E92A reveal a lipid-mediated closing mechanism. **(A)** Open and closed structures as represented by the first principal component and projections of 10 1 μs MD started from the open and closed state along PC1. **(B)** PC1 projection for 10 1 µs of the WT MthK pore domain started from the open state. **(C)** PC1 projection for 10 1 µs of the E92A MthK pore domain started from the open state. (**D**) Non-proteinogenic Cryo-EM density in closed-state E92A MthK, likely from lipid tails. (**E**) Fraction of MD simulations that stay in the open state, defined as PC1 < 5. **(F)** 2D density plot of lipid heavy atom count in the pore cavity vs PC1 projection for 10 10 µs simulations of E92A. **(G)** 2D density plot of water molecule count in the pore cavity vs PC1 projection for 10 10 µs simulations of E92A. Red dots highlight coordinates of **(H)** representative snapshots of states found in closing trajectories of MthK E92A MD simulations. Only 3 of 4 subunits are shown; the front subunit is hidden for clarity. K^+^ ions are shown in purple, water molecules are represented by blue spheres and selected POPC molecules are shown in orange.

**Figure 4.**
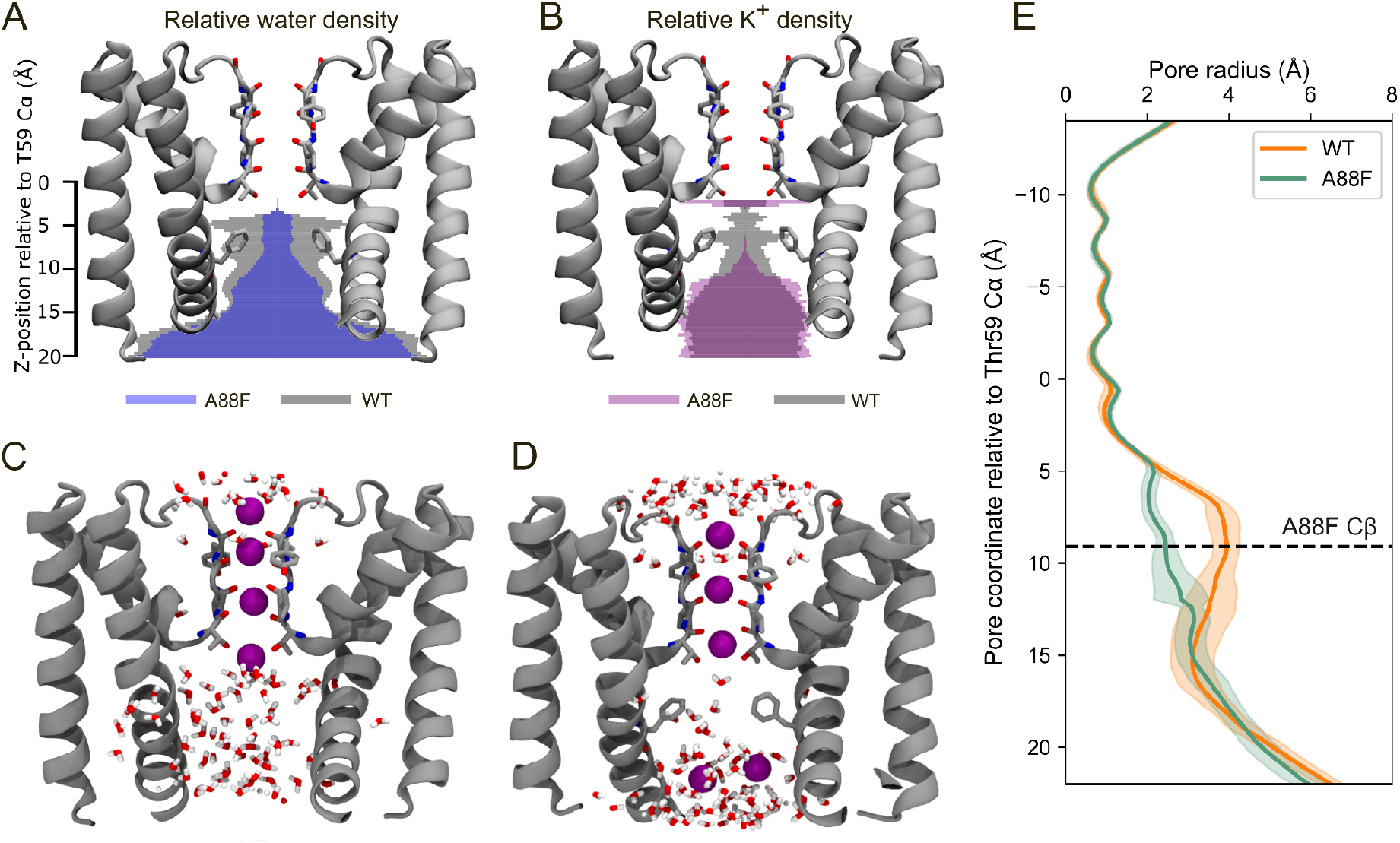
MthK A88F structure and blocking mechanism. **(A)** Simulated water density in the pore cavity relative to the S4 SF binding site in A88F mutant overlaid on the WT density. **(B)** Simulated K^+^ ion density in the pore cavity relative to the S4 SF binding site in A88F mutant overlaid on the WT density **(C)** Representative MD snapshot of K^+^ and water in the WT MthK cavity **(D)** Representative MD snapshot of K^+^ and water in the A88F MthK cavity. PHE88 side chains form an additional barrier for hydrated K^+^ along the permeation pathway. **(E)** Pore radius profiles along the ion conduction pathway from the last 200 ns of 10 replicates from pore domain simulations of WT and A88F MthK. Shaded area indicates standard deviation.

Interestingly, in our simulations of the full-length WT MthK channel, we did not observe major differences in conformational dynamics in simulations with and without Ca^2+^ ions. This is reflected in the behaviour of WT MthK pore domain - it samples conformations similar to the initial ones (open or closed). In contrast however, we observed several spontaneous closing events in simulations of the E92 mutant. Again, the conformations we sample are very similar for different membrane compositions (Fig. S4). It is somewhat surprising that we do see the conformational change in the E92A mutant but do not see it in the WT channel, even in the absence of calcium, since Ca^2+^ binding to the RCK is the main force driving gating of the WT channel. A reason for this could be that the larger conformational rearrangements required in the RCK domain and transmission of this signal to the pore is a slower process that we do not capture in our microsecond-scale MD simulations. In contrast the change in the pore domain is smaller and mainly concerns a rotation of the TM2 helix.^17^

### MD simulations of E92A capture a full open to closed transition

Capture of partial closings of E92A MthK in MD simulations of 1 μs length prompted us to perform extended simulations for both the WT and E92A mutant. We tracked the fraction of replicates that stayed below different thresholds along the open-closed transition (Fig. S6).

Using the approximate mid-point of PC1 = 5 nm as a cut-off, we see that all WT channels stay open for the full duration of the simulation (Fig. 3E, Fig. S6). In contrast, for E92A the majority of the channels close. With all 10 replicates of the pore domain reaching the halfway-threshold within 5 μs and 8 out of 10 full length channels reach this point within the 2.5 μs of simulation time. Closing rates are similar between the pore domain and the full-length channel.

Our success in obtaining several replicates that cover the full closing allows us to investigate in detail the molecular events occurring in this gating process. We find that there is a linear relationship between the degree of closure, as defined by PC1, and the number of lipid atoms present in the cavity. In the open state the cavity is lipid-free, in contrast to the closed state (Fig. 3F). The same analysis for water molecules yielded a wide range of hydration levels available near the open state, while intermediate and closed states are almost completely dry (Fig. 3G). The same trends are observed in the 2.5 μs simulations of the full length E92A mutant, where there is an initial dehydration of the cavity followed by collapse and lipid entry (Fig. S7C-D). Even in 10 μs simulations of the WT channel we do not observe substantial dewetting or lipid entry (Fig. S7A-B).

Visual inspection of trajectories that completely close, as defined by PC1 > 9 nm, reveal a consistent closing pathway (Fig. 3H, Fig. 6, Fig. S9, Movie 1). Importantly, this mechanism is the same in the full-length and pore-only channels, further validating the use of the pore domain as a model to study gating of the MthK channel. Starting from a 4-fold symmetric, hydrated, open state (Fig. 3H-I), the pore first collapses into a 2-fold symmetric dehydrated state. In this state two opposing helices move into the cavity, while the other two stay near their original positions (Fig. 3H-II). From here, two fenestrations open, through which lipid tails enter the pore and fill the cavity (Fig. 3H-III). In the last step, two more fenestrations open and two additional lipid tails enter and fill the cavity (Fig. 3H-IV). We believe this mechanism is also representative of the WT mechanism, as the end (open and closed) states for the mutants are identical to those resolved for the WT channel using CryoEM microscopy, including the presence of lipids in the upper cavity. Therefore, the main difference between WT and E92A is the lower energy barrier for initial narrowing of the lower gate and dehydration of the pore. The E92A mutant facilitates an initial narrowing of the lower gate, which leads to complete dehydration, whereas in the WT this process likely has a higher barrier necessitating an additional driving force from Ca^2+^ in the RCK domain transmitted by the linkers to the lower gate, that results in initial twist of the TM2 helices that creates a more hydrophobic cavity.

In support of lipid molecules entering the cavity, we resolved a non-protein density in the closed-state cavity of E92A MthK (Fig. 3D). This likely corresponds to lipid tails, which have previously been shown to be positioned near the cavity in cryo-EM structures of MthK as well as enter the cavity through fenestrations in MD simulations^15^.

### Structural mechanism for lower conductance in A88F

Stopped-flow assays of A88F MthK reveal greatly reduced flux compared to the WT channel, although higher than that of the E92A mutant (Fig. 1F). Previous electrophysiology recordings have also shown that while the open probability of A88F MthK is similar to that of WT, its single channel current amplitude is drastically reduced compared to both the WT and E92A MthK channels (Fig. 2B)^29,30^. This suggests that while both of these mutations result in a more hydrophobic cavity and reduced bulk activity, they do so via different mechanisms. We therefore also performed MD simulations of the open state A88F mutant pore. While we previously reported a cryo-EM structure of closed Ca^2+^-free A88F MthK(PDB:8FZ7),^17^ we were not successful in determining an experimental structure of Ca^2+^-bound A88F MthK. Thus, to simulate the A88F open channel, we mutated Ala88 to Phe *in silico* on the WT MthK open structure (PDB:3LDC)^29^. In simulations restrained to the open state the simulated single channel current is reduced ∼20 fold, in good qualitative agreement with the experimentally observed reduction in single channel current^29^.

In 10 1 µs simulations of the unrestrained A88F pore, no closing of the channel is observed. We see an open state very similar to the WT open state which is slightly narrower than the WT open state (Fig. S9). Simulations with position restraints reveal that the open states of WT and E92A are equally hydrated, with around 50 water molecules in the cavity, while only ∼30 water molecules reside in the A88F cavity (Fig. S10). This suggests that it is not the overall hydrophobicity or water content of the filter that leads to closing of the channel, but rather that the electrostatic and hydrophilic balance of residue 92 plays a key role in the regulation of channel closing.

To understand the large reduction in single-channel current amplitude in A88F MthK, we looked closely at the water and ion distributions in the cavity. In the WT pore, the cavity is completely hydrated and K^+^ ions can reach the S4 binding site of the SF in a solvated state (Fig. 5). In contrast, in A88F, the Phe88 side chains point towards the cavity and form a constriction, which also results in partial dewetting of the upper cavity (Fig. 5A, Fig. 5D, Fig. 5E), thereby resulting in (partial) desolvation of K^+^ ions on their way to the selectivity filter. This forms a large energy barrier for K^+^ ions along the permeation pathway and explains the much lower conductance of this channel. This is likely a very similar constriction mechanism to NaK2K, where the reverse equivalent mutant F92A is often used to increase conductance^32–34^. While in A88F the ion density in the lower cavity is unaffected, it is drastically reduced near the A88F mutation site and the profile lacks the secondary density peak below the Scavity binding site present in the WT channel (Fig. 5B).

**Figure 5.**
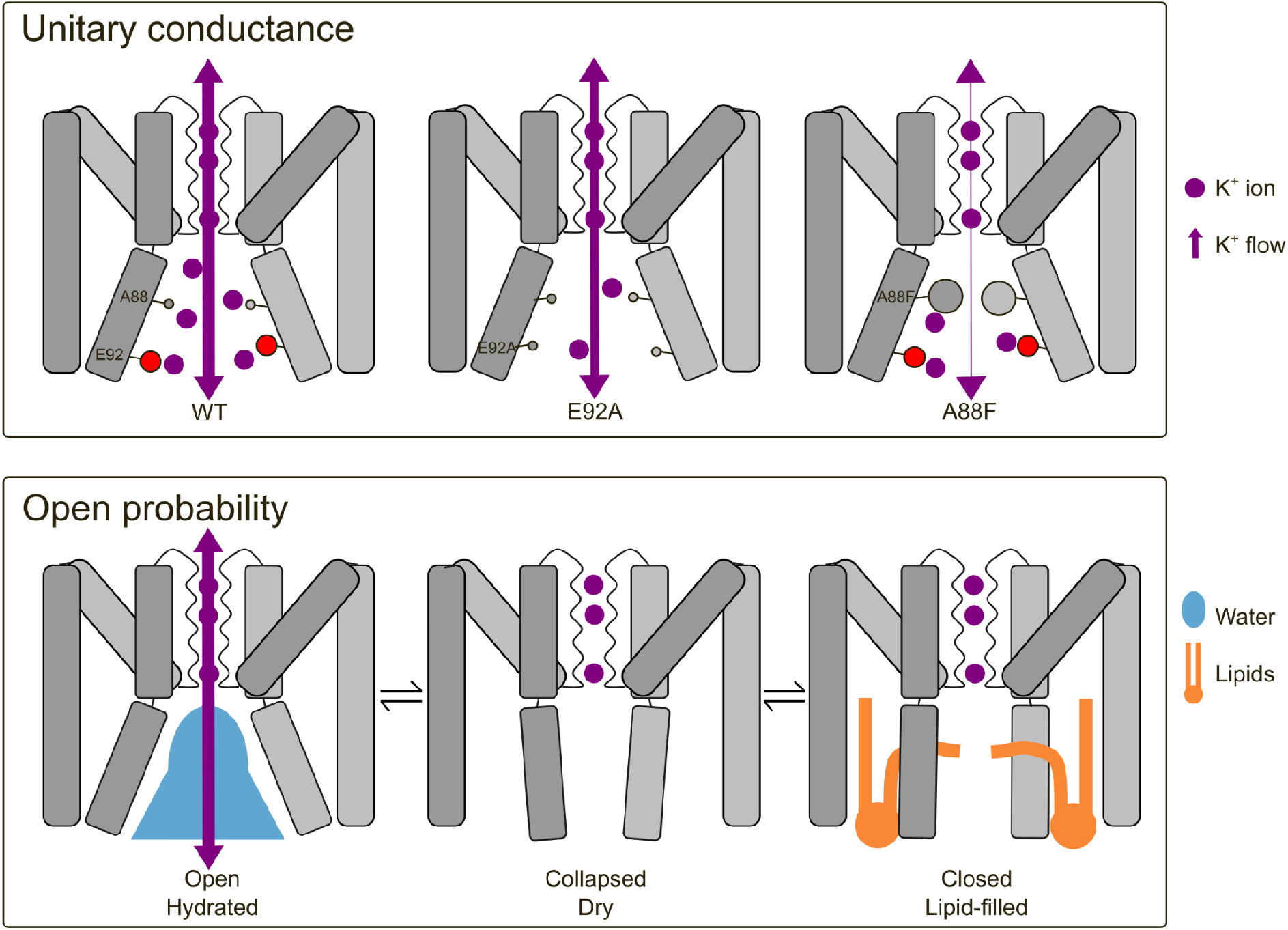
Summary. **Top**: mechanistic explanation of two different mechanisms for reduced single channel current in mutant channels. In E92A removal of the negatively charged glutamates leads to a lower effective K^+^ concentration in the cavity, which in turn leads to a lower conductance. In A88F a hydrophobic barrier is introduced, requiring (partial) desolvation of K^+^ ions. **Bottom:** Proposed lipid-mediated gating mechanism in MthK. Dehydration of the channel leads to an asymmetrically collapsed state. Lipids entering through fenestration can push this to a 4-fold symmetric canonical closed state.

### Determinants of channel conductance and gating

We investigated the molecular mechanism of reduced conductance in two MthK mutants. Although the A88F mutant and the E92A mutant are similar in nature, as they both increase the hydrophobicity of the MthK cavity, they modulate channel activity via two completely different mechanisms. In the A88F mutant we do not observe a change in the open-closed equilibrium, even though the upper cavity is dehydrated compared to the WT. The E92A mutation, lower in the cavity, however, does lead to closing of the channel. This is interesting as it highlights that it is not the overall hydrophobicity of the pore cavity that determines channel gating, but rather that residues in specific positions can be key determinants of channel gating. The removal of the glutamate residue in the E92A does not just make the cavity more hydrophobic, it also removes the negative charge. The electrostatic repulsion between these glutamates is likely another factor that contributes to the high open probability of the WT MthK channel. What is particularly interesting here is that the glutamate at this position (residue 92) is a unique feature of MthK.^35,36^ Other channels generally lack a negative charge at this position. For example, the equivalent residue is ALA108 in KcsA, where this has also been identified as a key residue of channel gating. The slightly more hydrophilic A108S and A108T mutations result in a moderate increase in channel activity and open probability.^37,38^ The A108D and A108E mutants both introduce a negative charge in this position, and this leads to drastic, pH dependent, increase in single channel current and open probability,^36^ indicating that this is indeed a key residue governing the permeation and gating of K^+^ channels.

### Comparison of MthK and BK gating

As MthK is often used as a model for BK channels^8,12,13^, and gating in both of these is facilitated by lipids^15,21,22^, we discuss the different gating mechanisms in a bit more detail. We find that the initial step in the open-to-closed transition of the MthK channel is dehydration of the cavity, which is subsequently followed by a rearrangement of the TM2 helices. Only when the cavity is dehydrated do lipids enter the cavity through the membrane-facing fenestrations and drive the channel into the canonical closed state with a four-fold symmetric helix bundle crossing (HBC).^15^ This differs from BK channels in several ways. Structures of non-conductive BK channels lack a HBC and the cavity is seemingly still solvent-accessible in this state.^25,39^ The open state of MthK lacks membrane-facing fenestrations, a large conformational change is necessary for lipids to enter the cavity from the side. ^15,16^ In BK, however, small openings are present in the open state, only a minor widening is needed to open fenestrations for lipid entry.^40^ These fenestrations open up further in the non-conductive state.^25,39^ This allows lipid molecules to enter the cavity and simultaneously displace water molecules from without a large conformational change.^21^ This is not possible in MthK as initial dehydration of the cavity is required to trigger the conformational change that opens the membrane-facing fenestrations. Classical hydrophobic gating, including a vapor barrier preventing ion permeation, is well supported in BK.^20,23,24^ It is unlikely that MthK shares this gating mechanism, as similar cavity dehydration leads to collapse of the cavity. This is further supported by the fact that Ca^2+^-free BK channels lack a classical closed structure with a HBC blocking the permeation pathway,^25,39^ while in MthK Ca^2+^-free structures are closed with a HBC.

## Conclusions

We studied the gating transition in the calcium-activated potassium channel MthK using molecular dynamics simulations, cryo-Electron microscopy and functional assays. We resolved new structures of one MthK mutant that increase the hydrophobicity of the pore cavity, and find open and closed state states identical to those of the WT channel, indicating their suitability as a model to study the gating mechanism of the MthK channel.

MD simulations and functional electrophysiology provide a complete mechanistic explanation of the previously observed lowered single channel conductance of these channels, summarized in the top panel of Figure 5. The E92A mutant has slightly lower single channel conductance due to a lower effective K^+^ concentration in the cavity; the negatively charged glutamate sidechains serve as an electrostatic funnel to attract K^+^ ions to the cavity. The effect of the A88F mutant on single channel current is much more severe as this mutant results in dewetting of the upper cavity and introduces a hydrophobic barrier along the ion permeation pathway. The additionally required desolvation of K^+^ ions is highly unfavourable and leads to a drastic drop in single channel current.

While A88F does not result in a decreased open probability, E92A severely shifts the open-closed equilibrium and decreases the initial barrier in the open-closed transition, by facilitating narrowing of the lower gate. This allowed us to study the complete closing pathway of the MthK channel for the first time, summarized in the bottom panel of figure 5. We identified a new intermediate, dehydrated, non-conductive state of the channel along the open-closed transition pathway. We also show the tight coupling between closing of the channel and lipid entry into the cavity; lipid tails entering through fenestrations push the channel from the collapsed intermediate state to the symmetrical closed state.

## Methods

### Protein expression and purification

Expression of MthK mutant channels (E92A, E92A Δ2-17) in *E.coli*, purification and reconstitution into nanodiscs were performed as previously described^31^. Briefly, the MthK mutant constructs were cloned in pQE82a vectors and transformed into XL-1 Blue cells (Agilent). Cells were grown at 37°C in Luria-Bertani media supplemented with 200 µg/ml ampicillin to an optical density of ∼1.0 and induced with 400 µM IPTG (US Biological) for 3 h at 37°C. Cell pellets were resuspended in buffer A (100 mM KCl, 50 mM Tris pH 7.6 with 50 µg/ml DNase (Sigma), 1 µM leupeptin/pepstatin (Roche) and 0.17 mg/ml PMSF (Roche)) and lysed by a probe sonication at 60-75% power. Membrane proteins were extracted by incubation with 50 mM decyl maltoside (DM; Anatrace) for 2 h rotation at room temperature. The suspension was centrifuged at 17,500g for 45 min at 17°C and the supernatant was loaded on pre-equilibrated Co^2+^ column with buffer B (100 mM KCl, 50 mM Tris pH 7.6, 5 mM DM). After washing the column with 20 mM imidazole, the bound proteins were eluted with 400 mM imidazole. Thrombin (Roche) was added to cleave the 6xHis tag in room temperature for 3 h. Subsequently, the MthK proteins were further purified by size exclusion chromatography on a Superdex-200 column (GE Healthcare) in buffer B. Peak fractions were pooled and concentrated to 8-10 mg/ml for further experiments.

### Nanodisc reconstitution

MthK E92A-containing nanodiscs were assembled as described previously ^31,41^. Purified MthK protein was mixed with MSP1E3 (Addgene plasmid #20064) and a lipid mixture of POPE:POPG (3:1; Anatrace) at a molar ratio of 1:2:85 (MthK monomer:MSP1E3:lipid). Lipids were solubilized in buffer containing 100 mM KCl, 20 mM HEPES pH 7.5, and 2% (w/v) CHAPS. Detergent removal was initiated by the addition of Bio-Beads SM-2 (20 mg per 100 µl mixture; Bio-Rad) and incubation for 3 h at room temperature with gentle rotation, followed by a second incubation for 12 h. The supernatant was filtered through a 0.22 μm Spin-X centrifugation tube filter (Costar) and loaded onto a Superose 6 (GE Lifesciences) equilibrated with 100 mM KCl and 20 mM HEPES pH 8.5. Peak fractions were collected and concentrated to ∼10 mg/ml for cryo-EM analysis.

### Grid Preparation and cryo-EM data collection

The cryo-EM samples were prepared as previously described ^31,41^. MthK E92A nanodisc samples were supplemented with 3 mM Fos8-F, and 5 mM CaCl_2_ (for open state) or 1 mM EDTA (for closed state) immediately before grids freezing. For the open-state dataset, 4,174 micrographs were collected on a Titan Krios microscope equipped with a K2 direct electron detector (NYSBC Krios1) at a pixel size of 1.073 Å and a total electron dose of 68.15 e^−^ Å^−2^. For the closed-state dataset, 3,577 micrographs were collected on a Titan Krios equipped with a K3 detector (NYU Krios) at a pixel size of 0.852 Å and a total dose of 53.09 e^−^ Å^−2^.

### Cryo-EM data processing and model building

The cryo-EM datasets were processed as previously described^31^. The closed-state dataset was processed in cryoSPARC. For the open-state dataset, initial refinement was performed using cryoSPARC non-uniform refinement, followed by further processing in RELION 3.0. Atomic models for the closed state were refined in PHENIX using the WT closed-state structure as the initial model. Owing to the lower resolution of the open-state 1 maps, the model was rigid-body fitted into the density using UCSF Chimera not subjected to further refinement or PDB deposition.

### Reconstitution into liposomes

Reconstitution of MthK mutant channels into large unilamellar vesicles (LUVs) for functional assays was performed as previously described ^31,41,42^. Purified MthK mutant protein (450 µg) was mixed with 15 mg of 3:1 DOPC:POPG (Anatrace) solubilized by sonication in reconstitution buffer containing 100 mM KNO_3_, 10 mM HEPES pH 7.0, and 35 mM CHAPS. The mixture was incubated at room temperature for 30 min after addition of 1 ml of an ANTS-containing solution (100 mM KNO_3_, 10 mM HEPES pH 7.0, 75 mM ANTS (Life Technologies)). LUV formation was induced by the addition of 1 g of Bio-Beads SM-2 along with 1 ml of reconstitution buffer, followed by rotation for 3 h at room temperature. The solution became visibly turbid after Bio-Beads treatment, consistent with vesicle formation. Vesicles were subsequently extruded through a 100 nm polycarbonate membrane using a Mini-Extruder (Avanti Polar Lipids). Unencapsulated ANTS was removed using PD-10 desalting columns (GE Healthcare) equilibrated with assay buffer (140 mM KNO_3_, 10 mM HEPES pH 7.0). The desalting was performed following the manufacturer’s protocol: 2.5 ml of sample was loaded onto the column and eluted with 3 ml of assay buffer, and the eluted fraction was collected. The resulting vesicle suspension was diluted 10-fold with assay buffer prior to measurements to achieve an appropriate signal-to-noise ratio.

### Stopped-flow fluorescence flux assays and data analysis

Stopped-flow measurements of Tl^+^ flux were carried out using an SX-20 stopped-flow spectrophotometer (Applied Photophysics), as described previously ^42–44^. MthK-containing liposomes were mixed with a premix buffer (140 mM KNO_3_, 10 mM HEPES pH 11.7, 34.4 mM Ca(NO_3_)_2_ and incubated in a delay loop for defined time intervals. Subsequently, the sample was mixed with either a control buffer (140 mM KNO_3_, 10 mM HEPES pH 8.5, 17.2 mM Ca(NO_3_)_2_ or a quench buffer (94 mM KNO_3_, 50 mM TlNO_3_, 10 mM HEPES pH 8.5, 17.2 mM Ca(NO_3_)_2_), and ANTS fluorescence was recorded. For each experimental condition, measurements were repeated six to eight times. Fluorescence traces were analyzed using MATLAB (MathWorks). The initial 100 ms of the quench signal was then fitted with a stretched exponential function. Data represent the mean ± S.D. from three independent LUV preparations.

### MD Simulations

For simulations of the pore domain the same simulation box as in our previous work was used^35,45^. For the full length open-state channel we used the 6OLY structure^5^ and added missing N-termini for the RCK domain using PyMOL^46^. For both the full-length and pore-only simulations of the closed-state channel we used the 6U5R cryoEM structure^15^, where the RCK domain was manually removed. For the simulations of the N-type inactivated state the pore domain of the 6U68 structure was used. Structures were embedded in a POPC bilayer and a solvated box with ∼0.8 mM KCl using the CHARMM-GUI bilayer builder^47^. The box was equilibrated following the 6 step equilibration procedure provided by CHARMM-GUI where position restraints are gradually reduced in consecutive equilibration runs. For the A88F and E92A mutants mutations were added after equilibration using the PyMOL mutagenesis tool. For the simulations with POPE:POPG membranes the lipid converter python script^48^ was used to change the lipid composition. For simulations with Ca^2+^ we used a multisite calcium model^49^, gmx genion was used to add 10 Ca^2+^ ions (∼30 mM) and these were converted to the multisite model using the provided script. Systems were simulated using GROMACS 2022.6^50,51^. For simulations under voltage simulations an external electric field of 300 mV was applied along the Z-axis, based on the box size in the Z-dimension after equilibration. We performed 10 simulations of at least 1 μs each for all conditions we test. For the longer simulations we extended the full length channels simulations to 2.5 μs and the simulations of the pore domain of WT and E92A MthK to 10 μs. We used the CHARMM36m forcefield^52^ along with the Charmm-modified TIP3P water model. In all simulations we used a 2 fs time step and the Particle Mesh Ewald Method^53^ for the electrostatic interactions, using a cut-off distance of 1.2 nm. The force-switch method was used to turn Van der Waals interactions off from 1.0 to 1.2 nm. Semi-isotropic Parrinello-Rahman pressure coupling^54^ and the Nose-Hoover thermostat^55^ were used to keep the system at 1 bar and 323 K, respectively. Hydrogen involving bonds were constrained using the LINCS algorithm^56^. In the production runs with position restraints on backbone atoms force constants of 1000 kJ/mol/nm^2^ were used. Atom densities in the cavity were calculated using a modified version of the MDAnalysis-based^57,58^ python script used before^35^, where the cavity was defined as a cylinder with a radius of 1.3 nm and a minimum and maximum distance 0 and 1.5 nm relative to SF CA carbon atoms. Permeation events were counted as previously described^45^. For the PCA the structures were manually reordered such that chain order matches for all structures. Main chain atoms for residues 24-98 were used. The set of structures used can be found in supplementary table 2.

### Alchemical transitions

Alchemical free energy calculations were performed as described previously for similar charge-changing systems^59,60^ using the PMX software^61,62^. Single transitions were performed for each chain separately. 455 frames per chain were extracted from the last 450 ns of 5 500 ns equilibrium simulations of the WT channel for the open and inactivated conformation.

## Data availability

The maps and models have been deposited in the Electron Microscopy Data Bank (EMDB) and the Protein Data Bank (PDB). Accession codes are: MthK-E92A mutant in the closed state (EMD-76624, PDB: 12OE), and MthK-E92A mutant in the open state (EMD-76634, PDB: 12OH). Raw fluorescence traces are available from the corresponding authors upon request. Sample raw data traces are provided in the corresponding figures. Input files and for MD simulations and final coordinates are deposited on zenodo and can be found at 10.5281/zenodo.19708726.

## Author contributions

B.L.d.G., C.M.N. and W.K. conceived and supervised the project. R.d.V. performed and analysed molecular dynamics simulations. C. F. and H.-S.Y. performed experiments. C. F., C.M.N. and H.-S.Y. analysed the experiments. R.d.V. prepared the manuscript with contributions from B.L.d.G., C.M.N., H.-S.Y. and W.K.

## Acknowledgments

R.d.V. thanks Carter J. Wilson for his help with PMX and the free energy calculations. C.M.N thanks Shubhangi Agarwal for help with the early stage of this project and cryo-EM data collection. R.d.V., W.K. and B.L.d.G. acknowledge funding from the Leibniz Collaborative Excellence Project “Ion Selectivity and Conduction Mechanism of Cation Channels” K305/2020. This work was sponsored in part by the NIH (GM088352 to C.M.N).

## References

(1) Hille, B. Ionic Channels in Excitable Membranes. Current Problems and Biophysical Approaches. Biophys. J. 1978, 22 (2), 283–294. 10.1016/S0006-3495(78)85489-7.

(2) Littleton, J. T.; Ganetzky, B. Ion Channels and Synaptic Organization: Analysis of the Drosophila Genome. Neuron 2000, 26 (1), 35–43. 10.1016/s0896-6273(00)81135-6.

(3) Doyle, D. A.; Cabral, J. M.; Pfuetzner, R. A.; Kuo, A.; Gulbis, J. M.; Cohen, S. L.; Chait, B. T.; MacKinnon, R. The Structure of the Potassium Channel: Molecular Basis of K+ Conduction and Selectivity. Science 1998, 280 (5360), 69–77. 10.1126/science.280.5360.69.

(4) Miller, C. An Overview of the Potassium Channel Family. Genome Biol. 2000, 1 (4), reviews0004.1. 10.1186/gb-2000-1-4-reviews0004.

(5) Kopec, W.; Rothberg, B. S.; de Groot, B. L. Molecular Mechanism of a Potassium Channel Gating through Activation Gate-Selectivity Filter Coupling. Nat. Commun. 2019, 10 (1), 5366. 10.1038/s41467-019-13227-w.

(6) Antz, C.; Fakler, B. Fast Inactivation of Voltage-Gated K+ Channels: From Cartoon to Structure. Physiology 1998, 13 (4), 177–182. 10.1152/physiologyonline.1998.13.4.177.

(7) Posson, D. J.; McCoy, J. G.; Nimigean, C. M. The Voltage-Dependent Gate in MthK Potassium Channels Is Located at the Selectivity Filter. Nat. Struct. Mol. Biol. 2013, 20 (2), 159–166. 10.1038/nsmb.2473.

(8) Jiang, Y.; Lee, A.; Chen, J.; Cadene, M.; Chait, B. T.; MacKinnon, R. The Open Pore Conformation of Potassium Channels. Nature 2002, 417 (6888), 523–526. 10.1038/417523a.

(9) Hummer, G.; Rasaiah, J. C.; Noworyta, J. P. Water Conduction through the Hydrophobic Channel of a Carbon Nanotube. Nature 2001, 414 (6860), 188–190. 10.1038/35102535.

(10) Water in Nanopores and Biological Channels: A Molecular Simulation Perspective | Chemical Reviews. 10.1021/acs.chemrev.9b00830 (accessed 2025-09-12).

(11) Aryal, P.; Sansom, M. S. P.; Tucker, S. J. Hydrophobic Gating in Ion Channels. J. Mol. Biol. 2015, 427 (1), 121–130. 10.1016/j.jmb.2014.07.030.

(12) Zadek, B.; Nimigean, C. M. Calcium-Dependent Gating of MthK, a Prokaryotic Potassium Channel. J. Gen. Physiol. 2006, 127 (6), 673–685. 10.1085/jgp.200609534.

(13) Jiang, Y.; Lee, A.; Chen, J.; Cadene, M.; Chait, B. T.; MacKinnon, R. Crystal Structure and Mechanism of a Calcium-Gated Potassium Channel. Nature 2002, 417 (6888), 515–522. 10.1038/417515a.

(14) Scheuring, S. Forces and Energetics of the Canonical Tetrameric Cation Channel Gating. Proc. Natl. Acad. Sci. 2023, 120 (28), e2221616120. 10.1073/pnas.2221616120.

(15) Fan, C.; Sukomon, N.; Flood, E.; Rheinberger, J.; Allen, T. W.; Nimigean, C. M. Ball-and-Chain Inactivation in a Calcium-Gated Potassium Channel. Nature 2020, 580 (7802), 288–293. 10.1038/s41586-020-2116-0.

(16) Ye, S.; Li, Y.; Jiang, Y. Novel Insights into K+ Selectivity from High-Resolution Structures of an Open K+ Channel Pore. Nat. Struct. Mol. Biol. 2010, 17 (8), 1019–1023. 10.1038/nsmb.1865.

(17) Fan, C.; Flood, E.; Sukomon, N.; Agarwal, S.; Allen, T. W.; Nimigean, C. M. Calcium-Gated Potassium Channel Blockade via Membrane-Facing Fenestrations. Nat. Chem. Biol. 2023, 1–10. 10.1038/s41589-023-01406-2.

(18) Aryal, P.; Abd-Wahab, F.; Bucci, G.; Sansom, M. S. P.; Tucker, S. J. A Hydrophobic Barrier Deep within the Inner Pore of the TWIK-1 K2P Potassium Channel. Nat. Commun. 2014, 5 (1), 4377. 10.1038/ncomms5377.

(19) Natale, A. M.; Deal, P. E.; Minor, D. L. Structural Insights into the Mechanisms and Pharmacology of K2P Potassium Channels. J. Mol. Biol. 2021, 433 (17), 166995. 10.1016/j.jmb.2021.166995.

(20) Gu, R.-X.; de Groot, B. L. Central Cavity Dehydration as a Gating Mechanism of Potassium Channels. Nat. Commun. 2023, 14 (1), 2178. 10.1038/s41467-023-37531-8.

(21) Mironenko, A.; Groot, B. L. de; Kopec, W. Lipid Gating of BK Channels and Mechanism of Activation by Negatively Charged Lipids. bioRxiv May 30, 2025, p 2025.05.27.656279. 10.1101/2025.05.27.656279.

(22) Coronel, L.; Di Muccio, G.; Rothberg, B. S.; Giacomello, A.; Carnevale, V. Lipid-Mediated Hydrophobic Gating in the BK Potassium Channel. Nat. Commun. 2025, 16 (1), 7354. 10.1038/s41467-025-61638-9.

(23) Jia, Z.; Yazdani, M.; Zhang, G.; Cui, J.; Chen, J. Hydrophobic Gating in BK Channels. Nat. Commun. 2018, 9 (1), 3408. 10.1038/s41467-018-05970-3.

(24) Nordquist, E. B.; Jia, Z.; Chen, J. Inner Pore Hydration Free Energy Controls the Activation of Big Potassium Channels. Biophys. J. 2023, 122 (7), 1158–1167. 10.1016/j.bpj.2023.02.005.

(25) Tao, X.; MacKinnon, R. Molecular Structures of the Human Slo1 K+ Channel in Complex with Β4. eLife 2019, 8, e51409. 10.7554/eLife.51409.

(26) Hite, R. K.; Tao, X.; MacKinnon, R. Structural Basis for Gating the High-Conductance Ca2+-Activated K+ Channel. Nature 2017, 541 (7635), 52–57. 10.1038/nature20775.

(27) Crystal structure of full-length KcsA in its closed conformation | PNAS. https://www.pnas.org/doi/10.1073/pnas.0810663106 (accessed 2025-09-11).

(28) Rohaim, A.; Vermeulen, B. J. A.; Li, J.; Kümmerer, F.; Napoli, F.; Blachowicz, L.; Medeiros-Silva, J.; Roux, B.; Weingarth, M. A Distinct Mechanism of C-Type Inactivation in the Kv-like KcsA Mutant E71V. Nat. Commun. 2022, 13 (1), 1574. 10.1038/s41467-022-28866-9.

(29) Shi, N.; Zeng, W.; Ye, S.; Li, Y.; Jiang, Y. Crucial Points within the Pore as Determinants of K+ Channel Conductance and Gating. J. Mol. Biol. 2011, 411 (1), 27–35. 10.1016/j.jmb.2011.04.058.

(30) Parfenova, L. V.; Crane, B. M.; Rothberg, B. S. Modulation of MthK Potassium Channel Activity at the Intracellular Entrance to the Pore. J. Biol. Chem. 2006, 281 (30), 21131–21138. 10.1074/jbc.M603109200.

(31) Fan, C.; Sukomon, N.; Flood, E.; Rheinberger, J.; Allen, T. W.; Nimigean, C. M. Ball-and-Chain Inactivation in a Calcium-Gated Potassium Channel. Nature 2020, 580 (7802), 288–293. 10.1038/s41586-020-2116-0.

(32) Derebe, M. G.; Sauer, D. B.; Zeng, W.; Alam, A.; Shi, N.; Jiang, Y. Tuning the Ion Selectivity of Tetrameric Cation Channels by Changing the Number of Ion Binding Sites. Proc. Natl. Acad. Sci. 2011, 108 (2), 598–602. 10.1073/pnas.1013636108.

(33) Öster, C.; Tekwani Movellan, K.; Goold, B.; Hendriks, K.; Lange, S.; Becker, S.; de Groot, B. L.; Kopec, W.; Andreas, L. B.; Lange, A. Direct Detection of Bound Ammonium Ions in the Selectivity Filter of Ion Channels by Solid-State NMR. J. Am. Chem. Soc. 2022, 144 (9), 4147–4157. 10.1021/jacs.1c13247.

(34) Sauer, D. B.; Zeng, W.; Canty, J.; Lam, Y.; Jiang, Y. Sodium and Potassium Competition in Potassium-Selective and Non-Selective Channels. Nat. Commun. 2013, 4 (1), 2721. 10.1038/ncomms3721.

(35) Öster, C.; de Vries, R.; Li, J.; Qoraj, D.; Lange, S.; Shi, C.; Kopec, W.; L de Groot, B.; Lange, A. Atomistic Mechanism of Calcium-Mediated Inward Rectification of the MthK Potassium Channel by Solid-State NMR and MD Simulations. J. Am. Chem. Soc. 2025. 10.1021/jacs.5c16155.

(36) Nimigean, C. M.; Chappie, J. S.; Miller, C. Electrostatic Tuning of Ion Conductance in Potassium Channels. Biochemistry 2003, 42 (31), 9263–9268. 10.1021/bi0348720.

(37) Paynter, J. J.; Sarkies, P.; Andres-Enguix, I.; Tucker, S. J. Genetic Selection of Activatory Mutations in KcsA. Channels Austin Tex 2008, 2 (6), 413–418. 10.4161/chan.2.6.6874.

(38) Irizarry, S. N.; Kutluay, E.; Drews, G.; Hart, S. J.; Heginbotham, L. Opening the KcsA K+ Channel: Tryptophan Scanning and Complementation Analysis Lead to Mutants with Altered Gating. Biochemistry 2002, 41 (46), 13653–13662. 10.1021/bi026393r.

(39) Contreras, G. F.; Shen, R.; Latorre, R.; Perozo, E. Structural Basis of Voltage-Dependent Gating in BK Channels. Nat. Commun. 2025, 16 (1), 5846. 10.1038/s41467-025-60639-y.

(40) Tao, X.; Hite, R. K.; MacKinnon, R. Cryo-EM Structure of the Open High-Conductance Ca2+-Activated K+ Channel. Nature 2017, 541 (7635), 46–51. 10.1038/nature20608.

(41) Correlating Ion Channel Structure and Function. In Methods in Enzymology; Elsevier, 2021; Vol. 652, pp 3–30. 10.1016/bs.mie.2021.02.016.

(42) Posson, D. J.; Rusinova, R.; Andersen, O. S.; Nimigean, C. M. Stopped-Flow Fluorometric Ion Flux Assay for Ligand-Gated Ion Channel Studies. In Potassium Channels; Shyng, S.-L., Valiyaveetil, F. I., Whorton, M., Eds.; Methods in Molecular Biology; Springer New York: New York, NY, 2018; Vol. 1684, pp 223–235. 10.1007/978-1-4939-7362-0_17.

(43) Rusinova, R.; Kim, D. M.; Nimigean, C. M.; Andersen, O. S. Regulation of Ion Channel Function by the Host Lipid Bilayer Examined by a Stopped-Flow Spectrofluorometric Assay. Biophys. J. 2014, 106 (5), 1070–1078. 10.1016/j.bpj.2014.01.027.

(44) Ingólfsson, H. I.; Andersen, O. S. Screening for Small Molecules’ Bilayer-Modifying Potential Using a Gramicidin-Based Fluorescence Assay. ASSAY Drug Dev. Technol. 2010, 8 (4), 427–436. 10.1089/adt.2009.0250.

(45) Hui, C.; de Vries, R.; Kopec, W.; de Groot, B. L. Effective Polarization in Potassium Channel Simulations: Ion Conductance, Occupancy, Voltage Response, and Selectivity. Proc. Natl. Acad. Sci. 2025, 122 (21), e2423866122. 10.1073/pnas.2423866122.

(46) Schrödinger, LLC. The PyMOL Molecular Graphics System, Version 2.5.4, 2015.

(47) CHARMM-GUI Membrane Builder: Past, Current, and Future Developments and Applications. 10.1021/acs.jctc.2c01246.

(48) Larsson, P.; Kasson, P. M. Lipid Converter, A Framework for Lipid Manipulations in Molecular Dynamics Simulations. J. Membr. Biol. 2014, 247 (11), 1137–1140. 10.1007/s00232-014-9705-5.

(49) Zhang, A.; Yu, H.; Liu, C.; Song, C. The Ca2+ Permeation Mechanism of the Ryanodine Receptor Revealed by a Multi-Site Ion Model. Nat. Commun. 2020, 11 (1), 922. 10.1038/s41467-020-14573-w.

(50) Abraham, M. J.; Murtola, T.; Schulz, R.; Páll, S.; Smith, J. C.; Hess, B.; Lindahl, E. GROMACS: High Performance Molecular Simulations through Multi-Level Parallelism from Laptops to Supercomputers. SoftwareX 2015, 1–2, 19–25. 10.1016/j.softx.2015.06.001.

(51) Berendsen, H. J. C.; van der Spoel, D.; van Drunen, R. GROMACS: A Message-Passing Parallel Molecular Dynamics Implementation. Comput. Phys. Commun. 1995, 91 (1), 43–56. 10.1016/0010-4655(95)00042-E.

(52) Huang, J.; Rauscher, S.; Nawrocki, G.; Ran, T.; Feig, M.; de Groot, B. L.; Grubmüller, H.; MacKerell, A. D. CHARMM36m: An Improved Force Field for Folded and Intrinsically Disordered Proteins. Nat. Methods 2017, 14 (1), 71–73. 10.1038/nmeth.4067.

(53) Darden, T.; York, D.; Pedersen, L. Particle Mesh Ewald: An N⋅log(N) Method for Ewald Sums in Large Systems. J. Chem. Phys. 1993, 98 (12), 10089–10092. 10.1063/1.464397.

(54) Parrinello, M.; Rahman, A. Polymorphic Transitions in Single Crystals: A New Molecular Dynamics Method. J. Appl. Phys. 1981, 52 (12), 7182–7190. 10.1063/1.328693.

(55) Hoover, W. G. Canonical Dynamics: Equilibrium Phase-Space Distributions. Phys. Rev. A 1985, 31 (3), 1695–1697. 10.1103/PhysRevA.31.1695.

(56) Hess, B.; Bekker, H.; Berendsen, H. J. C.; Fraaije, J. G. E. M. LINCS: A Linear Constraint Solver for Molecular Simulations. J. Comput. Chem. 1997, 18 (12), 1463–1472. 10.1002/(SICI)1096-987X(199709)18:12<1463::AID-JCC4>3.0.CO;2-H.

(57) Michaud-Agrawal, N.; Denning, E. J.; Woolf, T. B.; Beckstein, O. MDAnalysis: A Toolkit for the Analysis of Molecular Dynamics Simulations. J. Comput. Chem. 2011, 32 (10), 2319–2327. 10.1002/jcc.21787.

(58) Gowers, R. J.; Linke, M.; Barnoud, J.; Reddy, T. J. E.; Melo, M. N.; Seyler, S. L.; Domanski, J.; Dotson, D. L.; Buchoux, S.; Kenney, I. M.; Beckstein, O. MDAnalysis: A Python Package for the Rapid Analysis of Molecular Dynamics Simulations; LA-UR-19-29136; Los Alamos National Laboratory (LANL), 2019. 10.25080/Majora-629e541a-00e.

(59) Wilson, C. J.; Karttunen, M.; de Groot, B. L.; Gapsys, V. Accurately Predicting Protein pKa Values Using Nonequilibrium Alchemy. J. Chem. Theory Comput. 2023, 19 (21), 7833–7845. 10.1021/acs.jctc.3c00721.

(60) Wilson, C. J.; Gapsys, V.; de Groot, B. L. Improving pKa Predictions with Reparameterized Force Fields and Free Energy Calculations. J. Chem. Theory Comput. 2025, 21 (8), 4095–4106. 10.1021/acs.jctc.5c00031.

(61) Gapsys, V.; Michielssens, S.; Seeliger, D.; de Groot, B. L. Pmx: Automated Protein Structure and Topology Generation for Alchemical Perturbations. J. Comput. Chem. 2015, 36 (5), 348–354. 10.1002/jcc.23804.

(62) Behera, S.; Wilson, C. J.; Schmidt, L.; Groot, B.L.de. Free Energy Simulations to Quantitatively Study Biomolecule Stability and Binding. ChemRxiv May 19, 2025. 10.26434/chemrxiv-2025-2zbd8.

