## Supporting information for "Hydrophobic and lipid-mediated gating mechanism revealed by low-conductance MthK mutants"

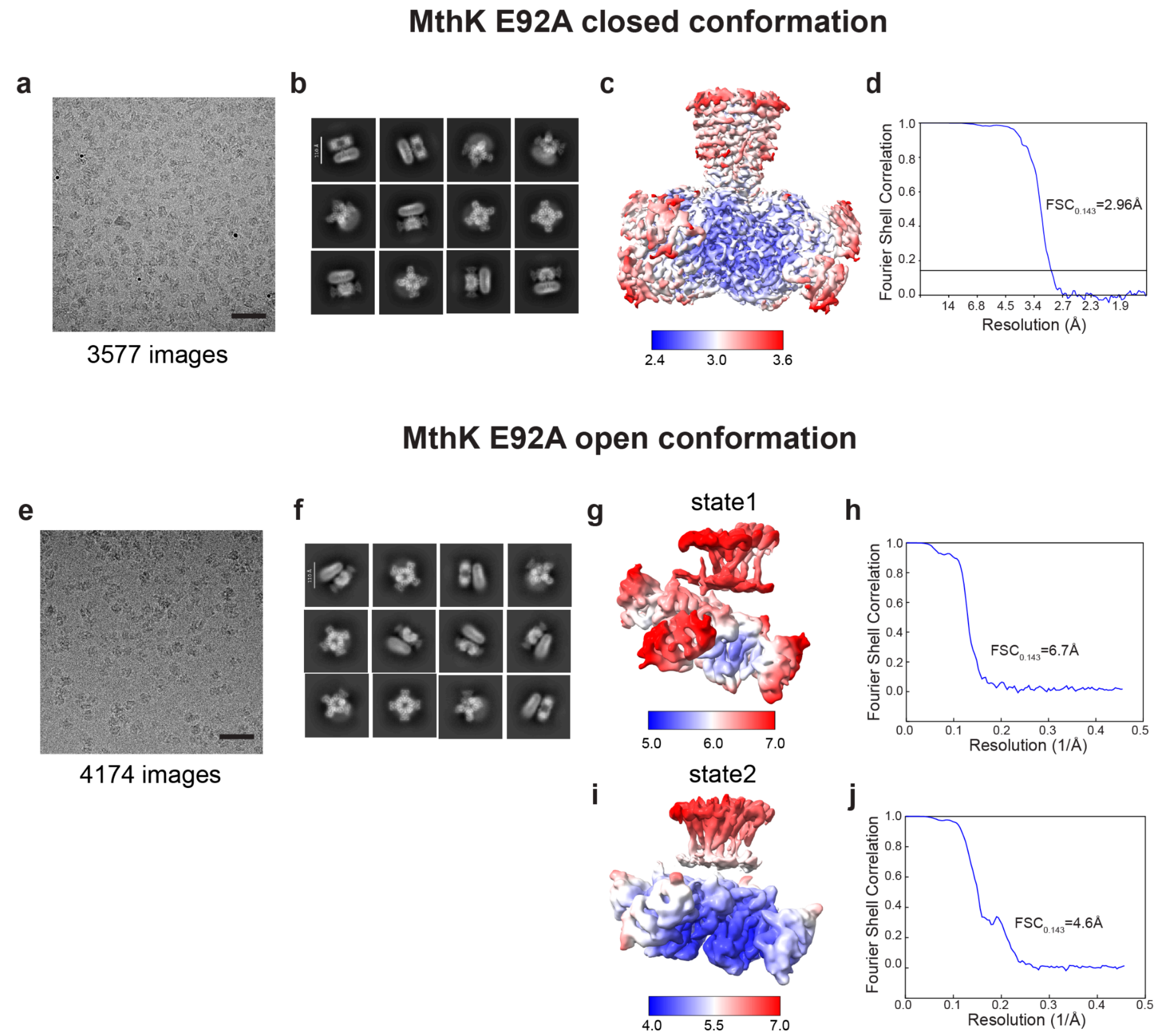


**Figure S1. Single-particle Cryo-EM characterization of closed and open MthK E92A mutants**. **a** Representative cryo-EM micrograph. Calibration bar is 40 nm. b-d Selected 2D class averages (**b**), local resolution maps colored in final cryo-EM map (**c**), and FSC curve (**d**) of MthK E92A closed conformation. **e** Representative cryo-EM micrograph. Calibration bar is 40 nm. f-h Selected 2D class averages (**f**), local resolution maps colored in final cryo-EM map (**g**), and FSC curve (**h**) of MthK E92A open conformation state 1. i-j Local resolution maps colored in final cryo-EM map (**i**), and FSC curve (**j**) of MthK E92A open conformation state 2.

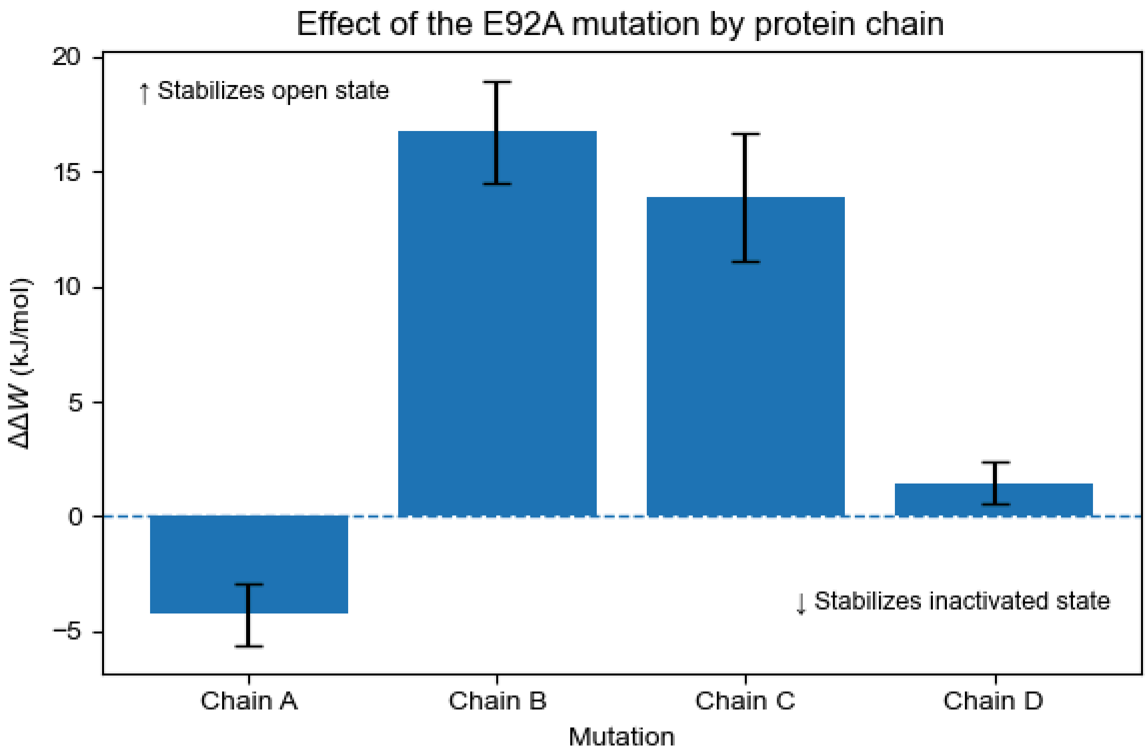


**Figure S2. The open state is favoured over the N-type inactivated state in the E92A mutant**

To determine whether the lower activity of the E92A mutant could be caused by an increase in N-type inactivation, we performed alchemical transitions from the E92 to E92A in each chain for both the open and the inactivated state. For each chain we computed the difference in nonequilibrium work between the open and inactivated states (ΔΔW) for the E92A mutation. Positive values indicate preferential stabilization of the activated state by the E92A mutation. Because simulations were performed in a single alchemical direction, we analyze differences in mean nonequilibrium work (ΔΔW) rather than equilibrium free energies; positive values indicate stabilization of the open state, negative values indicate stabilization of the inactivated state. We find that the E92A mutation stabilizes the open state over the inactivated state compared to the WT channel. Suggesting that the E92A mutation does not lead to an increase of N-type inactivation.

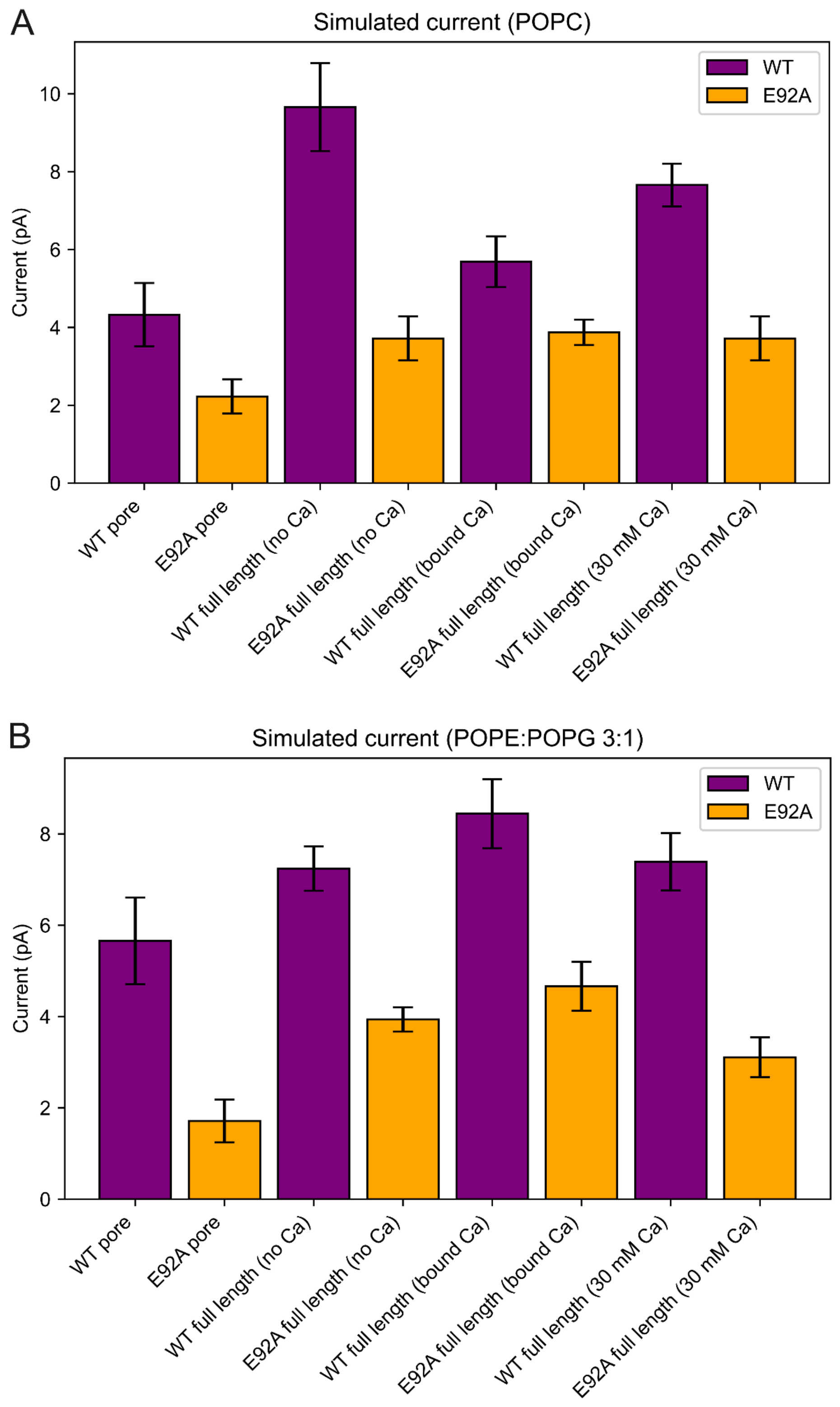


**Figure S3. Unitary conductance is reduced in the E92A mutant independent of membrane composition**

Simulated current for WT and E92A under 300 mV for the pore domain and the full length channel in the presence and absence of Ca in for different bilayer compositions. Average over 10 1 us replicas, error bars represent standard error of the mean. (**A**) Pure POPC bilayer (**B**) 3:1 POPE:POPG bilayer

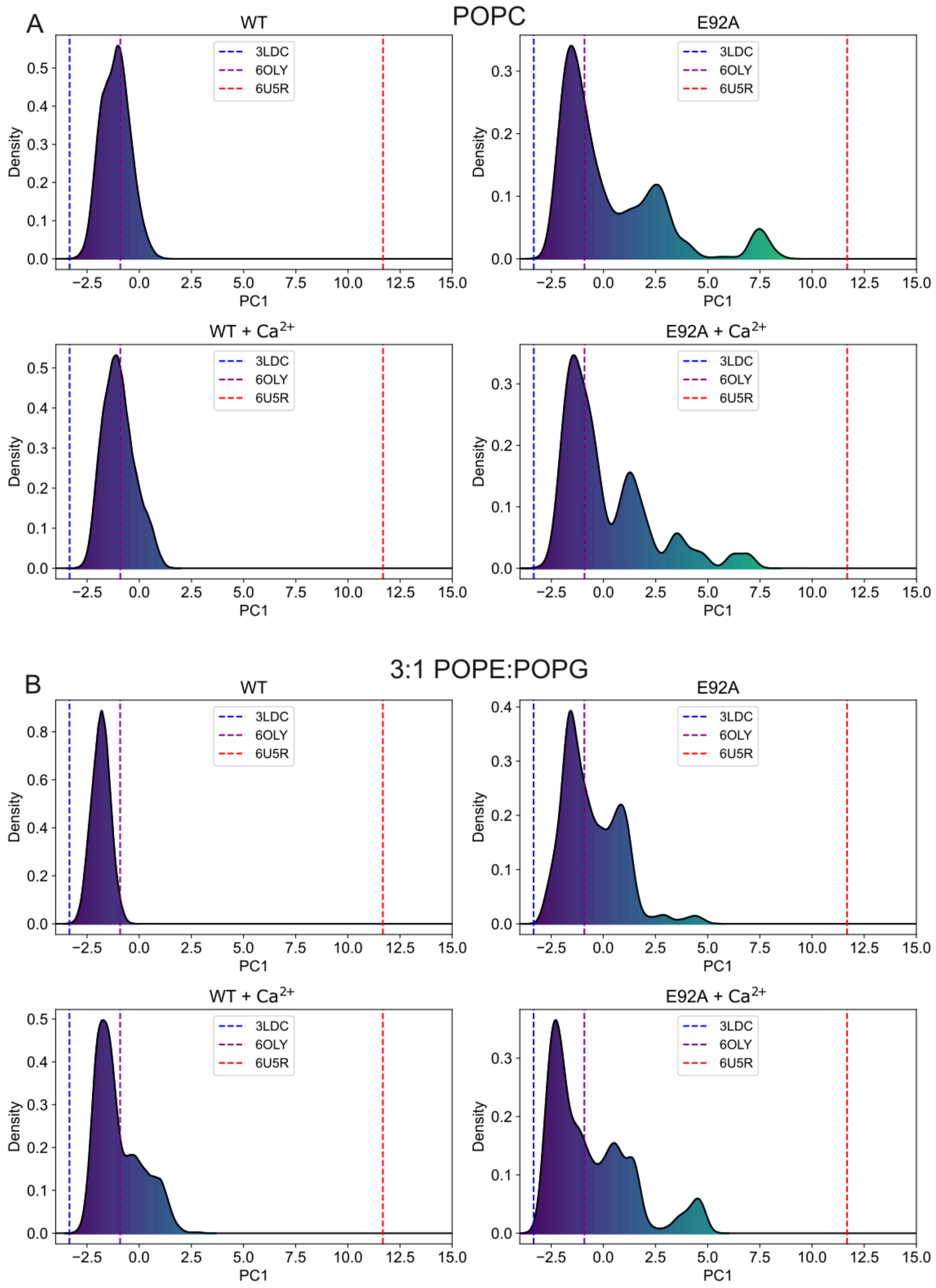


**Figure S4. Simulations of the E92A mutant sample more closed conformations independent of membrane composition**

PC1 projection for 10 1 µs of the full length WT and E92A channels in the absence and presence of $Ca^{2+}$ in different membrane compositions. (**A**) POPC (**B**) 3:1 POPE:POPG

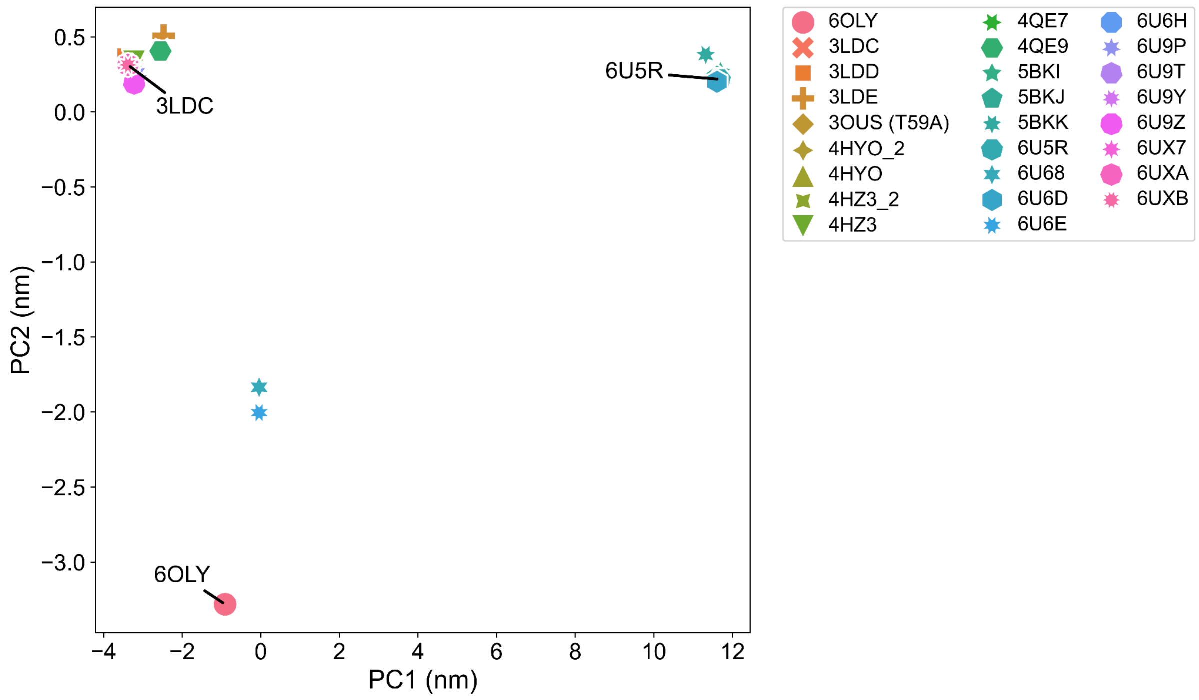


**Figure S5. 2D PCA projection of different experimentally resolved MthK structures.**

Separation along PC1 describes the open-closed transition.

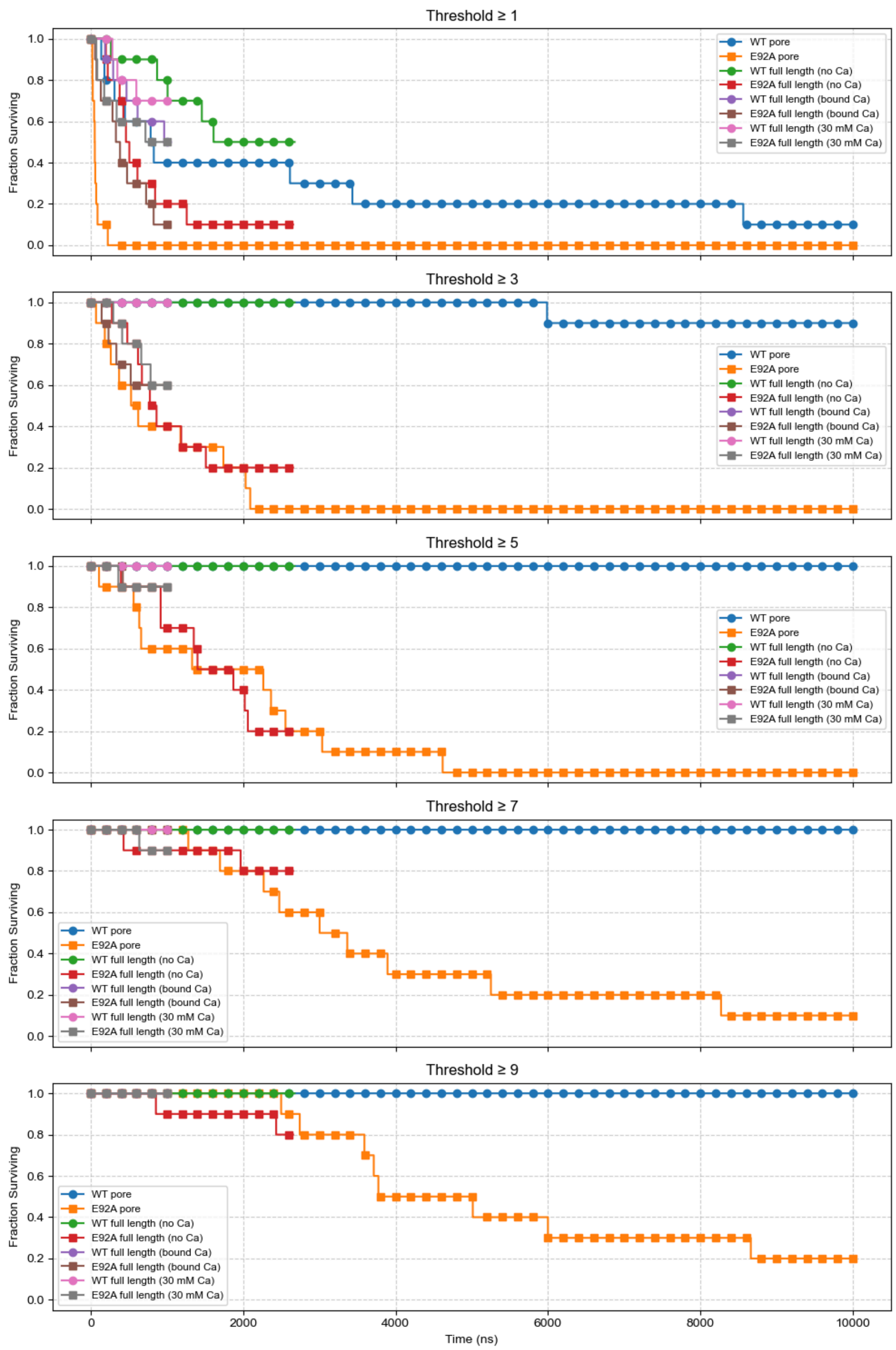


**Figure S6. The majority of E92A channels completely close within 10 µs**
Fraction of MD simulations that stay in the open state over time, using 5 different PC1 threshold values.

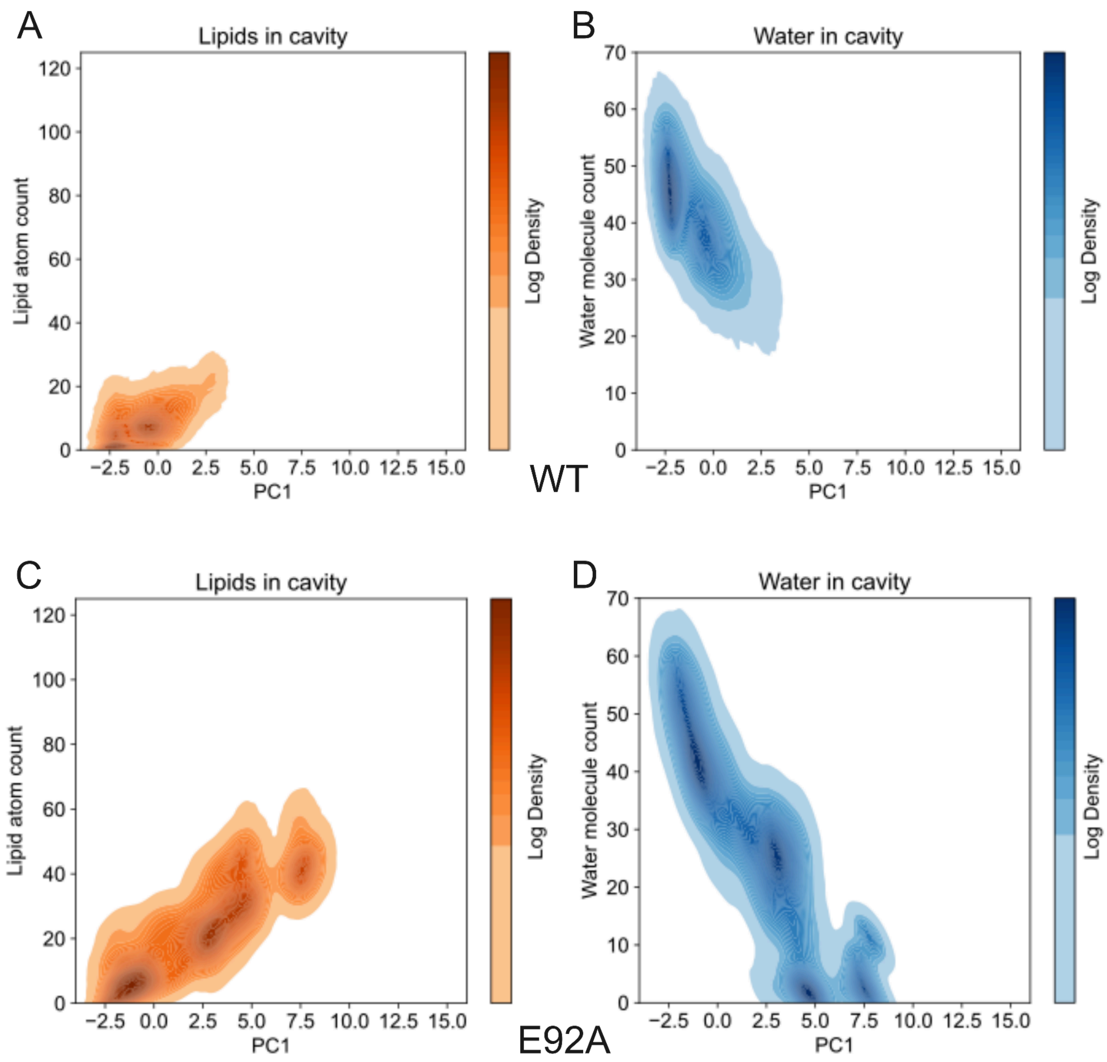


**Figure S7. Dehydration and lipid entry in full-length E92A follows the same trend as in the pore domain.**

**(A)** 2D density plot of lipid heavy atom count in the pore cavity vs PC1 projection for 10 10 µs simulations of the WT pore domain. **(B)** 2D density plot of water molecule count in the pore cavity vs PC1 projection for 10 2.5 µs simulations of the WT pore domain. **(C)** 2D density plot of lipid heavy atom count in the pore cavity vs PC1 projection for 10 2.5 µs simulations of full length E92A. **(D)** 2D density plot of water molecule count in the pore cavity vs PC1 projection for 10 2.5 µs simulations of full length E92A.

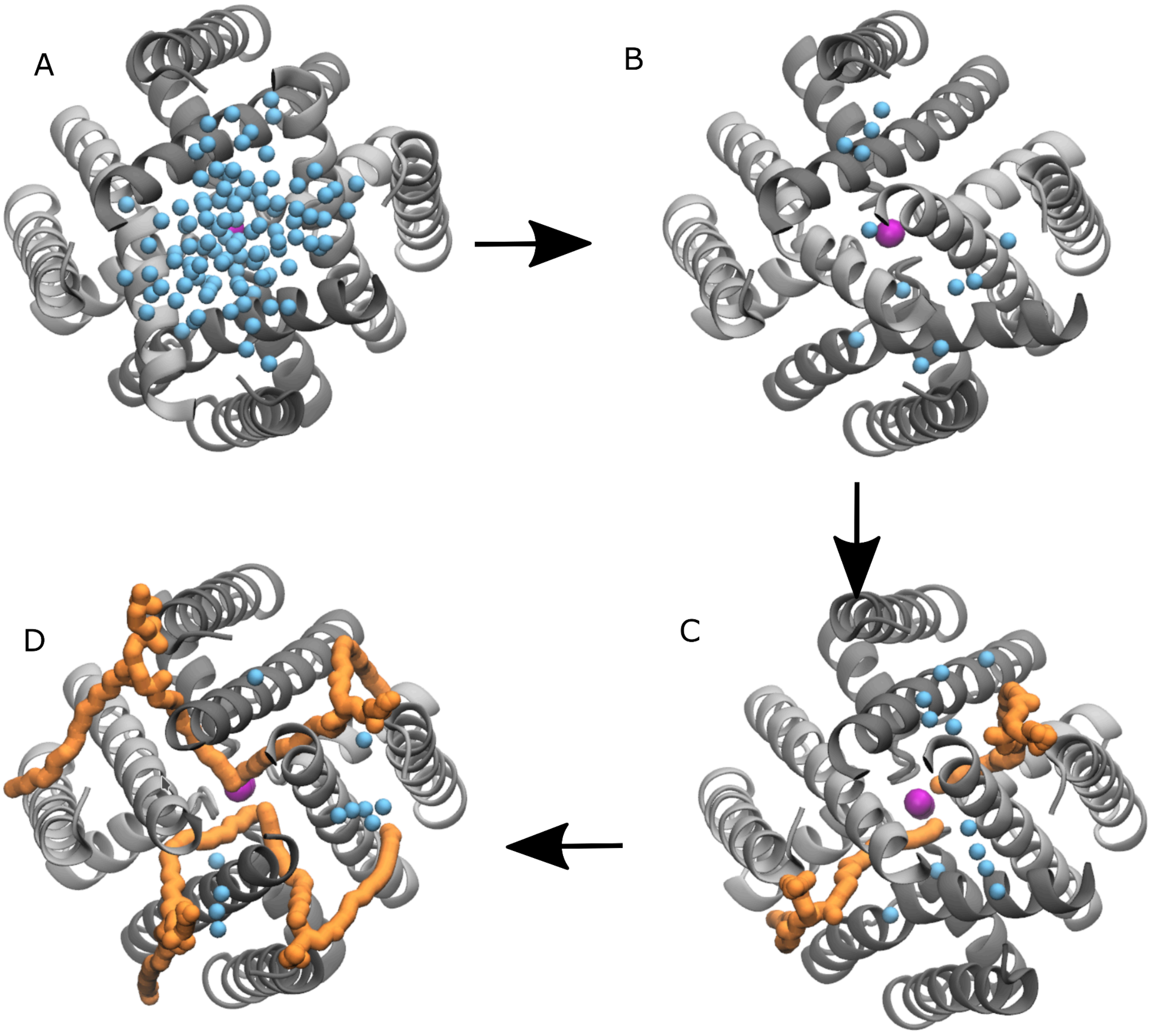


**Figure S8. Representative snapshots of states found in closing trajectories of MthK E92A MD simulations**

**(A)** 4-fold symmetric open state. The cavity is hydrated (blue beads) and lipid-free **(B)** 2-fold symmetric dehydrated state. The cavity dehydrates and 1 set of helices (in grey) moves closer together narrowing the cavity, while the other helices (in black) stay closer to the open configuration (**C**) Lipids enter the cavity through fenestrations from 2 sides. (**D**) Lipids now enter through all 4 fenestrations and the pore has reached the 4-fold symmetric closed state.

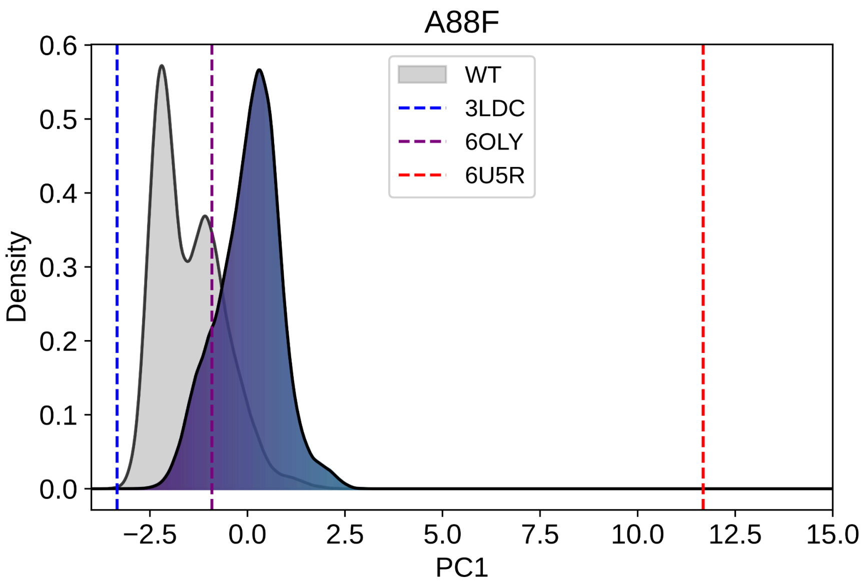


**Figure S9. The open A88F mutant is similar to the open WT channel**

PC1 projection for 10 1 µs of the A88F pore overlaid on the WT density.

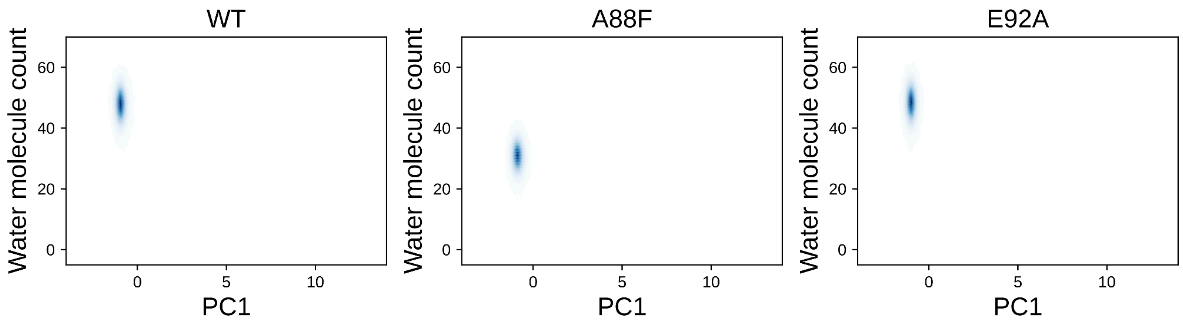


**Figure S10. The lower cavity is hydrated in open WT and mutant channels**

Number of water molecules in the cavity of WT, A88F and E92A pore domain MthK. Data from 10 1 µs simulations with the pore restrained to the open conformation.

**Table S1. Cryo-EM data collection, refinement, and validation statistics.**

| | MthK-E92A in open state | | MthK-E92A in closed state |
|---|---|---|---|
| | State 1 | State 2 (EMDB-76634) (PDB 12OH) | (EMDB-76624) (PDB 12OE) |
| **Data collection and processing** | | | |
| Magnification | 22500 | | 16500 |
| Voltage (kV) | 300 | | 300 |
| Electron exposure (e–/Å2) | 68.15 | | 53.09 |
| Defocus range (μm) | -1.4 to -2.6 | | -1.2 to -2.6 |
| Pixel size (Å) | 1.073 | | 0.852 |
| Symmetry imposed | C1 | | C4 |
| Final particles | 92,727 | 126,681 | 108,616 |
| Map resolution (Å) at FSC 0.143 | 6.7 | 4.6 | 2.96 |
| **Refinement** | | | |
| Initial model used (PDB code) | - | 3LDC | 6U6D |
| Model resolution (Å) at FSC 0.5 | - | 6.2 | 3.0 |
| Map sharpening B factor ($Å^2$) | - | -197 | -113.72 |
| Model composition | | | |
| Non-hydrogen atoms | - | 15980 | 16948 |
| Protein residues | - | 2104 | 2156 |
| Ligands | - | 0 | PEV:4 |
| B factor ($Å^2$) | | | |
| Protein | - | 203.1/630.37/307.40 | 2.37/131.46/50.23 |
| Ligand | - | - | 52.28/186.45/101.8 |
| R.m.s. deviations | | | |
| Bond lengths (Å) | - | 0.003 | 0.002 |
| Bond angles (º) | - | 0.700 | 0.444 |
| Validation | | | |
| MolProbity score | - | 1.88 | 1.45 |
| Clash score | - | 16.27 | 3.16 |
| Poor rotamer (%) | - | 0.06 | 2.24 |
| Ramachandran plot | | | |
| Favored (%) | - | 97.07 | 97.66 |
| Allowed (%) | - | 2.84 | 2.34 |
| Outliers (%) | - | 0.1 | 0 |

**Table S2. Structures used for the Principal Component Analysis**

| PDB ID | Method | State | Reference |
|---|---|---|---|
| 3ldc | X-RAY | Open | [1] |
| 3ldd | X-RAY | Open | [1] |
| 3lde | X-RAY | Open | [1] |
| 3ous | X-RAY | Open (T59A) | [2] |
| 3rbz | X-RAY | Open | [3] |
| 4hyo | X-RAY | Open | [4] |
| 4hz3 | X-RAY | Open | [4] |
| 4qe7 | X-RAY | Open | [5] |
| 4qe9 | X-RAY | Open | [5] |
| 5bki | CryoEM | Closed | [6] |
| 5bkj | CryoEM | Closed | [6] |
| 5bkk | CryoEM | Closed | [6] |
| 6oly | X-RAY | Open | [7] |
| 6u5r | CryoEM | Closed | [8] |
| 6u68 | CryoEM | Inactivated | [8] |
| 6u6d | CryoEM | Closed | [8] |
| 6u6e | CryoEM | Inactivated | [8] |
| 6u6h | CryoEM | Inactivated | [8] |
| 6u9p | X-RAY | Open | [9] |
| 6u9t | X-RAY | Open | [9] |
| 6u9y | X-RAY | Open | [9] |
| 6u9z | X-RAY | Open | [9] |
| 6ux7 | CryoEM | Open | [8] |
| 6uxa | CryoEM | Open | [8] |
| 6uxb | CryoEM | Open | [8] |

### References

(1) Ye, S.; Li, Y.; Jiang, Y. Novel Insights into K+ Selectivity from High-Resolution Structures of an Open K+ Channel Pore. *Nat. Struct. Mol. Biol.* **2010**, *17* (8), 1019–1023. https://doi.org/10.1038/nsmb.1865.

(2) Derebe, M. G.; Sauer, D. B.; Zeng, W.; Alam, A.; Shi, N.; Jiang, Y. Tuning the Ion Selectivity of Tetrameric Cation Channels by Changing the Number of Ion Binding Sites. *Proc. Natl. Acad. Sci.* **2011**, *108* (2), 598–602. https://doi.org/10.1073/pnas.1013636108.

(3) Pau, V. P. T.; Smith, F. J.; Taylor, A. B.; Parfenova, L. V.; Samakai, E.; Callaghan, M. M.; Abarca-Heidemann, K.; Hart, P. J.; Rothberg, B. S. Structure and Function of Multiple Ca2+-Binding Sites in a K+ Channel Regulator of K+ Conductance (RCK) Domain. *Proc. Natl. Acad. Sci.* **2011**, *108* (43), 17684–17689. https://doi.org/10.1073/pnas.1107229108.

(4) Posson, D. J.; McCoy, J. G.; Nimigean, C. M. The Voltage-Dependent Gate in MthK Potassium Channels Is Located at the Selectivity Filter. *Nat. Struct. Mol. Biol.* **2013**, *20* (2), 159–166. https://doi.org/10.1038/nsmb.2473.

(5) Guo, R.; Zeng, W.; Cui, H.; Chen, L.; Ye, S. Ionic Interactions of Ba2+ Blockades in the MthK K+ Channel. *J. Gen. Physiol.* **2014**, *144* (2), 193–200. https://doi.org/10.1085/jgp.201411192.

(6) Fan, C.; Flood, E.; Sukomon, N.; Agarwal, S.; Allen, T. W.; Nimigean, C. M. Calcium-Gated Potassium Channel Blockade via Membrane-Facing Fenestrations. *Nat. Chem. Biol.* **2023**, 1–10. https://doi.org/10.1038/s41589-023-01406-2.

(7) Kopec, W.; Rothberg, B. S.; de Groot, B. L. Molecular Mechanism of a Potassium Channel Gating through Activation Gate-Selectivity Filter Coupling. *Nat. Commun.* **2019**, *10* (1), 5366. https://doi.org/10.1038/s41467-019-13227-w.

(8) Fan, C.; Sukomon, N.; Flood, E.; Rheinberger, J.; Allen, T. W.; Nimigean, C. M. Ball-and-Chain Inactivation in a Calcium-Gated Potassium Channel. *Nature* **2020**, *580* (7802), 288–293. https://doi.org/10.1038/s41586-020-2116-0.

(9) Boiteux, C.; Posson, D. J.; Allen, T. W.; Nimigean, C. M. Selectivity Filter Ion Binding Affinity Determines Inactivation in a Potassium Channel. *Proc. Natl. Acad. Sci.* **2020**, *117* (47), 29968–29978. https://doi.org/10.1073/pnas.2009624117.
